# Optimistic reinforcement learning: computational and neural bases

**DOI:** 10.1101/038778

**Authors:** G. Lefebvre, M. Lebreton, F. Meyniel, S. Bourgeois-Gironde, S. Palminteri

## Abstract

While forming and updating beliefs about future life outcomes, people tend to consider good news and to disregard bad news. This tendency is supposed to support the optimism bias. Whether this learning bias is specific to “high-level” abstract belief update or a particular expression of a more general “low-level” reinforcement learning process is unknown. Here we report evidence in favor of the second hypothesis. In a simple instrumental learning task, participants incorporated better-than-expected outcomes at a higher rate compared to worse-than-expected ones. In addition, functional imaging indicated that inter-individual difference in the expression of optimistic update corresponds to enhanced prediction error signaling in the reward circuitry. Our results constitute a new step in the understanding of the genesis of optimism bias at the neurocomputational level.

## Introduction

"*It is the peculiar and perpetual error of the human understanding to be more moved and excited by affirmatives than negatives; whereas it ought properly to hold itself indifferently disposed towards both alike*" (p36.)^*^

People typically overestimate the likelihood of positive events and underestimate the likelihood of negative events. This cognitive trait in (healthy) humans is known as the optimism bias and has been repeatedly evidenced in many different guises and populations^1–3^ such as students projecting their salary after graduation^4^, women estimating their risk of getting breast cancer^5^ or heavy smokers assessing their risk of premature mortality^6^. One mechanism hypothesized to underlie this phenomenon is an asymmetry in belief updating, colloquially referred to as “the good news / bad news effect”^7,8^. Indeed, preferentially revising one’s beliefs when provided with favorable compared to unfavorable information constitutes a learning bias which could, in principle, generates and sustains an overestimation of the likelihood of desired events and a concomitant underestimation of the likelihood of undesired events (optimism bias)^9^.

This good news/bad news effect has recently been demonstrated in the case where outcomes are hypothetical future prospects associated with a strong a priori desirability or undesirability (estimation of post-graduation salary or the probability of getting cancer)^4,5^. In this experimental context, belief formation triggers complex interactions between episodic, affective and executive cognitive functions^7,8,10^, and belief updating relies on a learning process involving abstract probabilistic information^7,11–13^. However, it remains unclear whether this learning asymmetry also applies to immediate reinforcement events driving instrumental learning directed to affectively neutral options (i.e. with no a priori desirability or undesirability). If an asymmetric update is also found in a task involving neutral items and direct feedback, then the good news/bad news effect could be considered as a specific – cognitive – manifestation of a general reinforcement learning asymmetry. If the asymmetry were not found at the basic reinforcement learning level, this would mean that the asymmetry is specific to abstract belief updating, and this would require a theory explaining this discrepancy.

To arbitrate between these two alternative hypotheses, we fitted instrumental behavior of subjects performing a simple two-armed bandit task, involving neutral stimuli and actual and immediate monetary outcomes, with two learning models. The first model (a standard RL algorithm) confounded individual learning rates for positive and negative feedback and the second one differentiated them, potentially accounting for learning asymmetries.

Over two experiments, we found that subjects’ behavior was better explained by the asymmetric model, with an overall difference in learning rates consistent with preferential learning from positive, compared to negative, prediction errors.

Previous studies suggest that the good news/bad news effect is highly variable across subjects^11^. Behavioral differences in optimistic beliefs and optimistic update have been shown to be reflected by differences in brain activation in the prefrontal cortex^7^. However, the question remains whether or not and how such inter-individual behavioral differences are related to the inter-individual neural differences in the extensively documented reward circuitry^14^. Our imaging results indicate that the inter-individual variability in the tendency in optimistic learning correlates with prediction error-related signals in the reward system, including the striatum and the ventro-medial prefrontal cortex (vmPFC).

## Results

### Behavioral task and dependent variables

Healthy subjects performed a probabilistic instrumental learning task with monetary feedback, previously used in brain imaging, pharmacological and clinical studies^15–17^ (**Fig. 1A**). In this task, options (abstract cues) were presented in fixed pairs (i.e. conditions). In all conditions each cue was associated with a stationary probability of reward. In asymmetric conditions, the two reward probabilities differed between cues (25/75%). From asymmetric conditions we extracted the rate of “correct” response (selection of the best option) as a measure of performance (**Fig. 1B, left**). In symmetric conditions, both cues had the same reward probabilities (25/25% or 75/75%), such that there was no intrinsic “correct response”. In symmetric conditions we extracted for each subject and each symmetric pair, a “preferred response” rate, defined as the choice rate of the option most frequently selected by a given subject (i.e. by definition in more than 50% of trials). The preferred response rate, especially in the 25/25% condition, should be taken as a measure of the tendency to overestimate the value of one instrumental cue compared to the other, in absence of actual outcome-based evidence. (**Fig. 1B, right**). In a first experiment (N=50) that subjects performed while being fMRI scanned, the task involved reward (+0.5€) and reward omission (0.0€), as the best and worst outcome respectively. In a second purely behavioral experiment (N=35), the task involved reward (+0.5€) and punishment (-0.5€), as the best and worst outcome respectively. All the results presented in the main text concern Experiment 1, except those of the last section entitled "*Optimistic reinforcement learning is robust across different outcome valences*". Detailed behavioral and computational analyses concerning Experiment 2 are presented in **Supplementary Materials**.

**Fig. 1:**
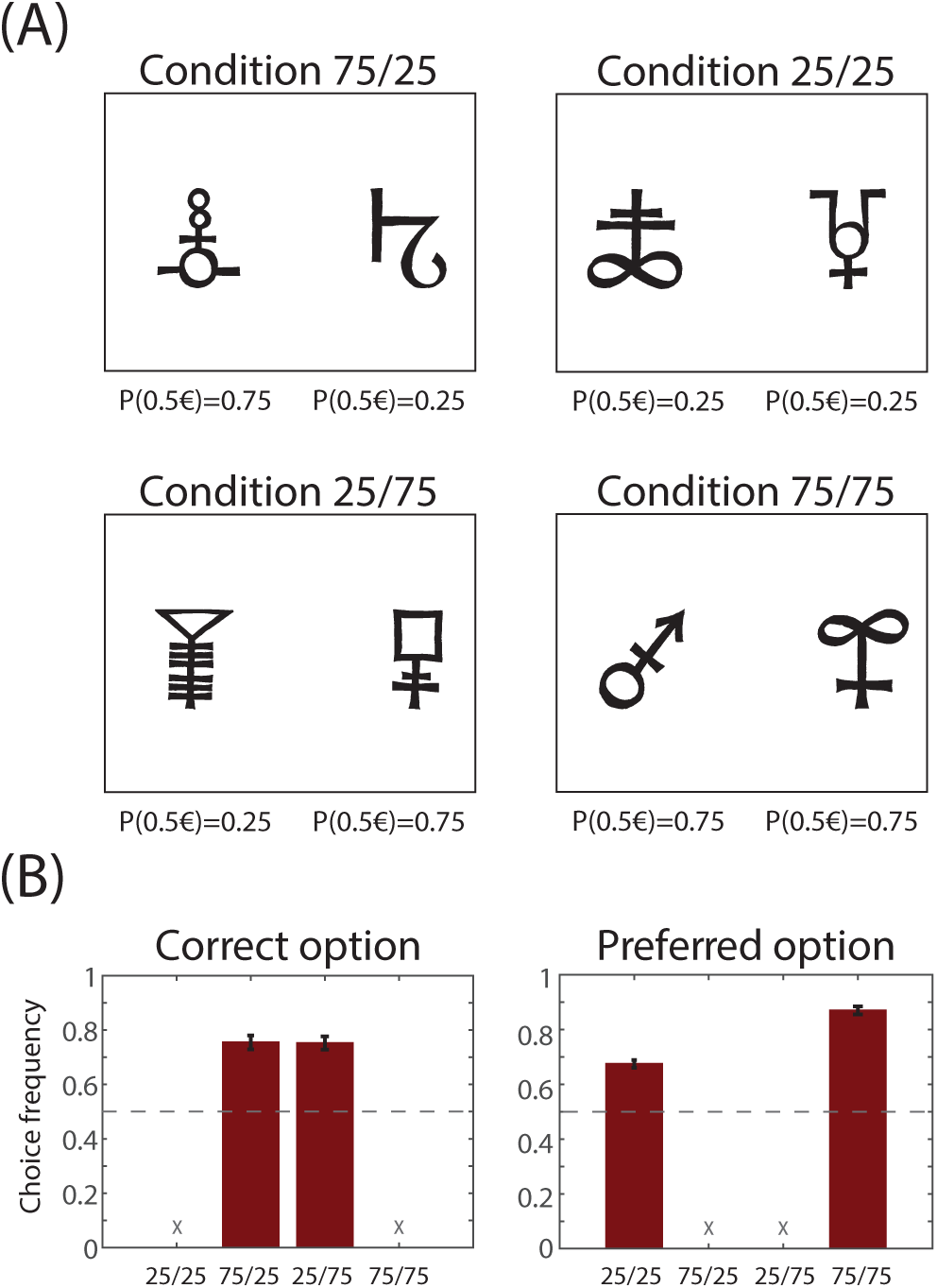
behavioral task and variables. **(A)** Task’s conditions and contingencies. Subjects selected between left and right symbols. Each symbol was associated with a stationary probability (*p* = 0.25 or 0.75) of winning 0.50€ and a reciprocal probability (1 – *p*) of getting nothing (first experiment) or losing 0.50€ (second experiment). In two conditions (rightmost column) the reward probability was the same for both symbols (“symmetric” conditions) and in two other conditions (leftmost column) the reward probability was different across symbols (“asymmetric” conditions). Note that the assignment between symbols conditions was randomized across subjects. (**B)** Dependent variables. In the leftmost panel, the histograms show the correct choice rate (i.e. choices directed toward the most rewarding stimulus in the asymmetric conditions). In the rightmost panel the histograms show the preferred option choice rate (i.e. the option chosen by subjects in more than 50% of the trials; this measure is relevant only in the symmetric conditions, where there is no intrinsic correct response). Bars indicate the mean and error bars indicate the SEM. Data are taken from both experiments (N=85).

### Computational models

We fitted the behavioral data with two reinforcement-learning models^18^. The “reference” model was represented by a standard Rescorla-Wagner model ^19^, thereafter referred to as RW model. The RW model learns option values by minimizing reward prediction errors. It uses a single learning rate (alpha: α) to learn from positive and negative prediction errors. The “target” model was represented by a modified version of the RW model, thereafter referred to as RW± model. In the RW± model, learning from positive and negative prediction errors is governed by different learning rates (alpha plus: α^+^ and alpha minus: α^-^ respectively). For α^+^ > α^-^ the RW± model instantiates optimistic reinforcement learning (i.e. the good news/bad news effect); for α^+^ = α^-^, the RW± instantiates unbiased reinforcement learning, just as in the RW model (the RW model is thus nested in the RW± model); finally, for α^+^ < α^-^ the RW± instantiates pessimistic reinforcement learning. In both models the choices are taken by feeding the option values into a softmax decision rule, whose exploration/exploitation trade-off is governed by a “temperature” parameter (β).

### Model comparison and model parameters analysis

We implemented Bayesian model comparison to establish which model better accounted for the behavioral data. For each model we estimated the optimal free parameters by maximizing the likelihood of the participants’ choices, given the models and sets of parameters. For each model and each subject, we calculated the Bayesian Information Criterion (BIC) by penalizing the maximum likelihood with the number of free parameters in the model. Random-effects BIC analysis indicated that the RW± model better explains the behavioral data compared to the RW model (BIC_RW_=99.4±4.4, BIC_RW±_=93.6±4.7; t(49)= 2.9, p=0.006, paired t-test), even accounting for its additional degree of freedom. A similar result was obtained when calculating the model exceedance probability using the BIC as an approximation of the model evidence^20^ (**Table 1**). RW± being the best fitting model we compared the learning rates fitted for positive (good news: α^+^ and negative (bad news: α^-^ prediction errors. We found α^+^ significantly higher compared to α^-^ (α^+^=0.36±0.05, α^-^=0.22±0.05, t(49)= 3.8, p<0.001 paired t-test). To summarize, model comparison indicated that, in our simple instrumental learning task, the best fitting model is the model with different learning rates for learning from positive and negative predictions errors (RW±). Crucially, learning rates comparison indicated that instrumental values are preferentially updated following positive prediction errors, which is consistent with an optimistic bias operating when learning from immediate feedback (optimistic reinforcement learning).

**Table 1.** models fitting and parameters in the two experiments. The table summarizes for each model its fitting performances and its average parameters: LLmax: maximal Log Likelihood; BIC: Bayesian Information Criterion (computed from LLmax); XP: exceedance probability; MF: model Frequency; α: learning rate for both positive and negative prediction errors (RW model); α^+^: learning rate for positive prediction errors; α^-^: average learning rate for negative prediction errors (RW± model); 1/β: average inverse of model temperature. Data are expressed as mean ± s.e.m. *P<0.01 comparing between the two models. #P<0.001 comparing between the two learning rates.

| Experiment / Model | LLmax | BIC | XP | MF | $\alpha$ | $\alpha^+$ | $\alpha^-$ | $1/\beta$ |
| --- | --- | --- | --- | --- | --- | --- | --- | --- |
| **Experiment 1 (N=50)** |  |  |  |  |  |  |  |  |
| RW Model | 45.1±2.2 | 99.4±4.4 | 0.17 | 0.43 | 0.32±0.05 | - | - | 0.16±0.03 |
| RW± Model | 40.0±2.4 | 93.6±4.7* | 0.83 | 0.57 | - | 0.36±0.05# | 0.22±0.05 | 0.13±0.03 |
| **Experiment 2 (N=35)** |  |  |  |  |  |  |  |  |
| RW Model | 44.2±2.9 | 97.6±5.9 | 0.10 | 0.39 | 0.24±0.05 | - | - | 0.53±0.16 |
| RW± Model | 38.1±3.0 | 89.8±6.0* | 0.90 | 0.61 | - | 0.45±0.06# | 0.18±0.05 | 0.30±0.10 |

### Computational phenotyping

To categorize subjects, we computed for each individual the between-model BIC difference (∆BIC=BIC_RW_ - BIC_RW±_) (see **Methods**). The ∆BIC quantifies at the individual level the goodness of fit improvement moving from the RW to the RW± model, or in other terms the fit improvement assuming different learning rates for positive and negative prediction errors. Subjects with a negative ∆BIC (N=25, in the first experiment) are subjects whose behavior is better explained by the RW model and therefore learn in an unbiased manner (thereafter refereed as RW subjects) (**Fig. 2A**). Subjects with a positive ∆BIC (N=25, in the first experiment) are subjects whose behavior is better explained by asymmetric learning (thereafter refereed as RW± subjects).

**Fig. 2:**
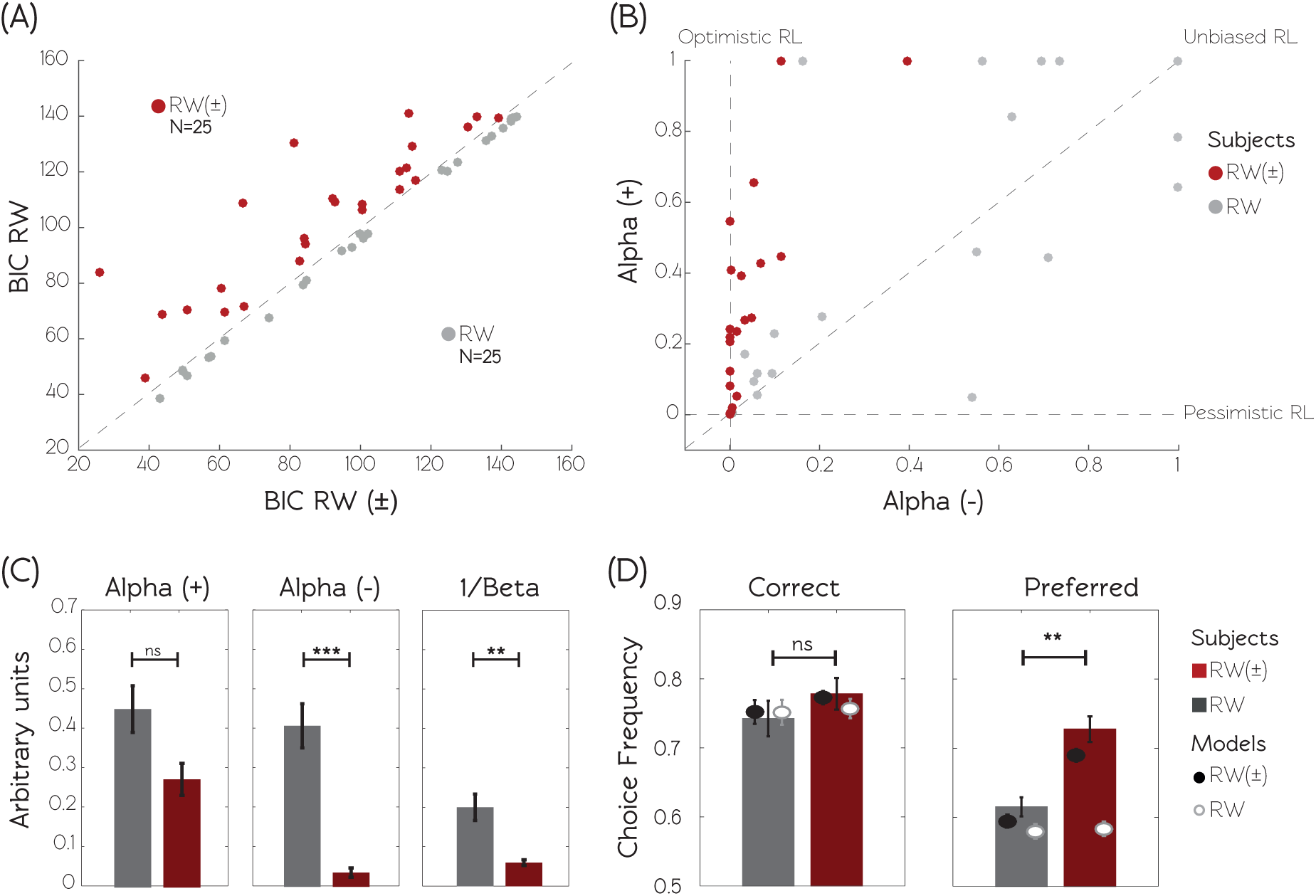
behavioral and computational identification of optimistic reinforcement learning. **(A)** Model comparison. The graphic displays the scatter plot of the BIC calculated for the RW model as a function of the BIC calculated for the RW± model. Smaller BIC values indicate better fits. Subjects are clustered in two populations according to the BIC difference (∆BIC = BIC_RW_ - BIC_RW±_) between the two models. RW± subjects (displayed in red) are characterized by a positive ∆BIC, indicating that the RW± model better explains their behavior. RW subjects (displayed in grey) are characterized by a negative ∆BIC, indicating that the RW model better explains their behavior. (**B)** Model parameters. The graphic displays the scatter plot of the learning rate following positive prediction errors (*α*^+^) as a function of the learning rate following negative prediction errors (*α*^-^), obtained from the RW± model. “Standard” reinforcement learners are characterized by similar learning rates for both types of prediction errors. “Optimistic” learners are characterized by a bigger learning rate for positive compared to negative prediction errors. “Pessimistic” learners are characterized by the opposite pattern. **C** The histograms show the RW± model free parameters (the learning rates + and - and the inverse temperature 1/β) as function of the subjects’ populations. **D** Actual and simulated choice rates. Histograms represent the observed and dots represent the model simulations of choices for both populations and both models, respectively for correct option (extracted from asymmetric condition), and from preferred option (extracted from the symmetrical condition 25/25%, see **Fig. 1A**). Model simulations are obtained using the individual best fitting free parameters. *p<0.05, ** p<0.01, ***p<0.001, two-sample two-sided t-test. Data are taken from the first experiment (N=50).

To test this hypothesis, learning rates fitted with the RW± model were entered in a two-way ANOVA with group (RW and RW±) and learning rates type (α^+^ and α^-^ as respectively between- and within-subjects factors. The ANOVA showed a main effect of learning rate type (F(1,48)=16.5, P<0.001) with α^+^ higher than α^-^. We also found a main effect of group (F(1,48)=10.48, P=0.002) and a significant group x learning type interaction (F(1,48)= 7.8, p=0.007). Post-hoc tests revealed that average learning rates for positive prediction errors were not different among the two groups, α^+^_RW_=0.45 ± 0.08 and α^+^_RW±_= 0.27 ± 0.06 (t(48) = 1.7, p=0.086, two-sample t-test). On the contrary, average learning rates for negative prediction errors were significantly different between groups, α^-^_RW_= 0.41 ± 0.08 and α^-^ _RW±_= 0.04 ± 0.02 (t(48)= 4.6, p<0.001, two-sample t-test). In addition, an asymmetry in learning rates was detected within the RW± group, where α^+^ was higher than α^-^ (t(24)=5.1, p<0.001, paired t-test) but not within RW group (t(24)=0.9, p=0.399, paired t-test). Thus, RW± subjects specifically drove the learning rates asymmetry found in the whole population. On the other side the RW subjects display “unbiased” (as opposed to “optimistic”) instrumental learning (**Fig. 2B and 2C**).

Interestingly, the exploration rate (captured by the 1/β, “temperature” parameter) was also found to be significantly different between the two groups of subject, 1/β_RW_=0.20 ± 0.05 and 1/β_RW±_=0.06 ± 0.01. (t(48)= 2.9, p=0.006, two-sample t-test). Importantly, the maximum likelihood of reference model (RW) was not different between the two groups of subjects, indicating similar baseline quality of fit (94.94±5.00 and 103.91±3.72 for RW and RW± subjects respectively, t(48)= -1.0, p=0.314, two-sample t-test). Accordingly the difference in the exploration rate parameter cannot be explained by difference in the quality of fit (i.e. noisiness of the data). This suggests that optimistic reinforcement learning, observed in RW± subjects, is also associated with exploitative, as opposite to explorative, behavior (**Fig. 2C**). Importantly, model simulations-based assessment of parameters recovery indicated that the two effects (learning rate asymmetry and lower exploration/exploitation trade-off) can be independently and correctly retrieved, ruling out the possibility that this twofold result is an artifact of the parameter optimization procedure (see **Supplementary Materials** and **Fig. S7**). To summarize, RW± subjects tend to weight more positive feedback and, as a consequence, to exploit more consistently the previously rewarded options (optimism).

### Behavioral signature distinguishing optimistic from unbiased subjects

In order to analyze the behavioral consequences of optimistic, as opposed to unbiased, learning and to confirm our model-based results with model-free behavioral observations, we compared the task’s dependent variables between our two groups of subjects (**Fig. 2D, Table 2**). Correct response rate did not differ between groups (t(48)=-0.7323, p=0.467, two-sample t-tests). However, the preferred response rate in the 25/25% condition was significantly higher for RW± group in comparison to RW group (t(48)= -3.4, p=0.001, two-sample t-test). Note that the same analysis performed on the 75/75% condition provided similar results (t(48)= -2.66, p=0.01, two-sample t-test).

**Table 2.** Behavioral and simulated data. The table summarizes for each experiment and each group of subjects, behavioral and simulated dependent variables: both real and simulated Correct Response in asymmetric conditions and both real and simulated Preferred Response in 25/25% condition. Data are expressed as mean ± s.e.m (in percentage). *P<0.01 two sample t-test.

| Experiment / Model | Correct Response | Correct Response (RW model) | Correct Response (RW ± model) | Preferred Response | Preferred Response (RW model) | Preferred Response (RW± model) |
| --- | --- | --- | --- | --- | --- | --- |
| Condition(s) | Asymmetric |  |  | Symmetric (25/25%) |  |  |
| Experiment 1 (N=50) |  |  |  |  |  |  |
| RW Group | 74.25 ± 3.65 | 75.20 ± 2.55 | 75.35 ± 2.42 | 61.5 ± 1.94 | 58.14 ± 0.67 | 59.47 ± 0.81 |
| RW± Group | 77.83 ± 3.25 | 75.58 ± 1.94 | 77.75 ± 1.44 | 72.75 ± 2.63* | 58.84 ± 0.55 | 69.36 ± 0.99 |
| Experiment 2 (N=35) |  |  |  |  |  |  |
| RW Group | 73.28± 4.63 | 73.65 ± 3.58 | 73.72± 3.49 | 61.89 ± 2.12 | 57.34 ± 1.14 | 59.82 ± 1.35 |
| RW± Group | 75.23± 4.73 | 75.29 ± 2.63 | 77.70 ± 1.86 | 73.73 ± 3.34* | 58.33 ± 0.98 | 70.74 ± 2.08 |

In order to validate the ability of RW± model to capture this difference, we performed simulations using both models and submitted them to the same statistical analysis as actual choices (**Fig. 2D**). The RW± model simulated preferred response rate was significantly higher for RW± group compared to the RW group (25/25%: t(48)= -5.4496, p<0.001; 75/75%: t(48)=-2.2670, p= 0.028; two-sample t-tests), which replicated human behavior. However, the simulated preferred response rates from the RW model were similar in the two groups (t(48)=0.566, p=0.566; 75/75%: t(48)=0.7448, p=0.4600; two-sample t-test), which departed from our observations in real subjects. This effect was particularly interesting in poorly rewarding environment (25/25%), where optimistic subjects tend to overestimate the value of one of the two options (**Fig. S1**). Finally, the preferred response rate in the symmetric conditions significantly correlated with both the computational features distinguishing RW and RW± subjects (normalized learning rates asymmetry (α^+^ - α^-^) / (α^+^ + α^-^): R=-0.475, P<0.001; choice randomness 1/β: R=-0.630, P<0.001). The preferred response rate thus provides a model-free signature of optimistic reinforcement learning that is congruent with our model simulation analysis: the preferred response rate was higher in RW± group in comparison to RW group and only simulations realized with RW± model were able to replicate this pattern of responses.

### fMRI signatures distinguishing optimistic from unbiased subjects

To investigate the neural correlates of the computational differences between RW± and RW subjects, we analyzed the brain activity both at the decision and outcome moments, using functional Magnetic Resonance Imagining (fMRI) and a model-based fMRI approach^21^. We devised a general linear model in which we modeled as separated events the choice and the outcome onset, each modulated by different parametric modulators. In a given trial, the choice onset was modulated by the chosen option Q-value (*Q*_*Chosen*_(*t*)), and the outcome onset was modulated by the reward prediction error (*δ*(*t*)). Concerning the choice onset, we found a neural network including the dmPFC and anterior Insulae negatively encoding *Q*_*Chosen*_(*t*), (P_FWE_<0.05 with a minimum of 60 continuous voxels) (**Fig. 3A and 3B**) (**Table 3**). We then tested for between-group differences within these two regions and found no significant difference (dmPFC: t(48)=0.0985, P=0.9220; Insulae t(48)=-0.0190, P=0.9849; two-sample t-tests) (**Fig. 3C**). Concerning the outcome onset we found a neural network including the striatum and vmPFC positively encoding δ(*t*), (P_FWE_<0.05 with a minimum of 60 continuous voxels) (**Fig. 3A and 3B**) (**Table3**). We then tested for between-group differences within these two regions and found significant differences (Striatum: t(48)=-3.2769, P=0.0020; vmPFC t(48)=-2.2590, P=0.0285; two-sample t-tests) (**Fig. 3C**). It therefore seems that the behavioral difference we observed between RW and RW± subjects finds its counterpart in a differential outcome-related signal in the ventral striatum. Within the regions displaying a between-group difference, we looked for correlation with the two computational features distinguishing optimistic from unbiased subjects. Interestingly, we found a positive and significant correlation between the striatal and vmPFC δ(*t*)-related activity and the normalized difference between learning rates (Striatum: R=0.4324, p=0.0017; vmPFC: R=0.3238,p=0.0218), but no significant difference between the same activity and 1/β (Striatum: R=-0.130, p=0.366; vmPFC: R=-0.272,p=0.3665), which suggests a specific link between this neural signature and the optimistic update.

**Fig. 3:**
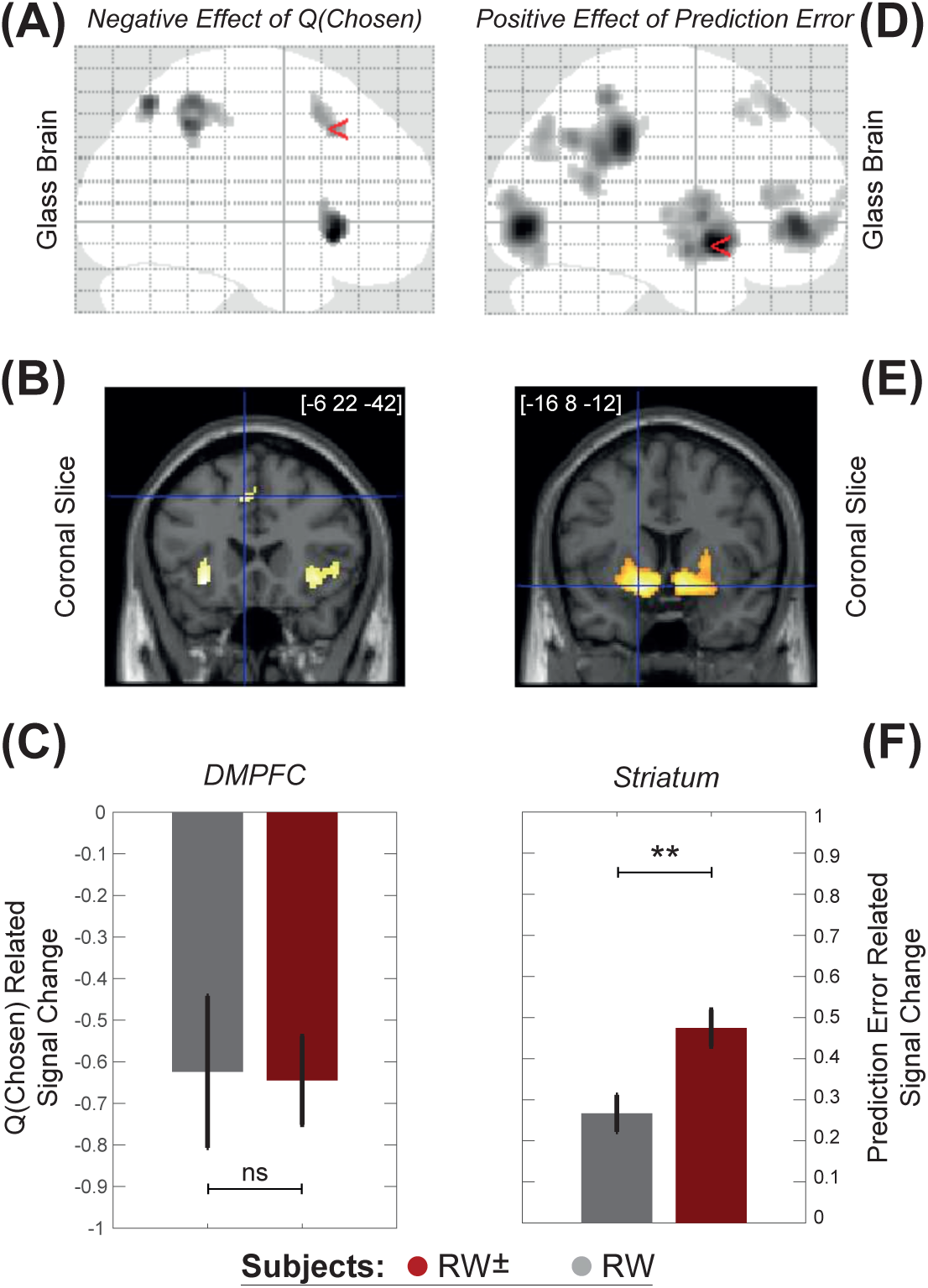
Functional signatures of the optimistic reinforcement learning. **(A)** and **(B)** Choice correlation. Statistical parametric maps of BOLD signal negatively correlating with the Q_Chosen_(t) at the choice onset. Areas colored in gray-to-black gradient on the axial glass brain and red-to-white gradient on the coronal slice show a significant effect (*p<0.001 corrected*). (**C**) Inter-individual differences. Histogram shows Q_Chosen_(t)-related signal change in DMPFC at the time of choice onset for both populations. Bars indicate the mean and error bars indicate the SEM. *p<0.05, unpaired t-tests. Data are taken from the first experiment (N=50). [*x, y, z*] coordinates are given in the MNI space (**D**) and (**F**) Outcome correlation. Statistical parametric maps of BOLD signal positively correlating with δ(t) at the outcome onset. Areas colored in gray-to-black gradient on the axial glass brain and red-to-white gradient on the coronal slice show a significant effect (*p<0.001 corrected*). (**F**) Inter-individual differences. Histogram shows δ(t)-related signal change in the striatum at the time of reward onset for both populations. Bars indicate the mean and error bars indicate the SEM. *p<0.05, **p<0.01 unpaired t-tests. Data are taken from the first experiment (N=50). [*x, y, z*] coordinates are given in the MNI space

**Table 3.** Activation table. FWE<0.05 whole brain corrected and 60 minimum voxels.

| Variable | Chosen option value (negative correlation) |  |  |
| --- | --- | --- | --- |
| Region (AAL) | Coordinates [x y z] | t value | Cluster size |
| Insula (left) | -30 22 -8 | 7.15 | 131 |
| Insula (right) | 34 24 -6 | 6.76 | 147 |
| superior parietal gyrus | -20 -66 54 | 6.56 | 112 |
| Angular gyrus | 40 -48 44 | 6.52 | 288 |
| superior frontal gyrus/Medial | -6 22 42 | 5.99 | 116 |
| Variable | Prediction error (positive correlation) |  |  |
| Region (AAL) | Coordinates [x y z] | t value | Cluster size |
| Putamen | -16 8 -12 | 11.02 | 1137 |
| Calcarine fissure | 2 -84 -4 | 10.7 | 1346 |
| Median cingulate | 0 -36 36 | 10.17 | 1533 |
| Caudate | 10 6 -10 | 10.15 | 984 |
| Anterior cingulate | -6 46 -4 | 9.56 | 911 |
| Angular gyrus | -50 -44 56 | 7.68 | 103 |
| superior frontal gyrus/dorsolateral | -18 38 52 | 6.7 | 167 |
| Angular gyrus | -40 -74 38 | 6.63 | 219 |
| Inferior frontal gyrus / triangular part | -44 32 10 | 6.54 | 165 |
| Cerebellum | 22 -76 -18 | 6.41 | 62 |

### Optimistic reinforcement learning is robust across different outcome valences

In the first experiment, getting nothing (0.0€) was the worst possible outcome. It could be argued that optimistic reinforcement learning (i.e. greater learning rate for positive than negative prediction errors: α^+^>α^-^) is dependent on the low negative motivational salience attributed to a neutral outcome and would not resist if negative prediction errors are accompanied by actual monetary losses. In order to confirm the independence of our results from outcome valence, in the second experiment the worst possible outcome was represented by a monetary loss (-0.5€), instead of reward omission (0.0€) as in the first experiment.

First, the second experiment replicated the model comparison result of the first experiment. Group-level BIC analysis indicated that the RW± model again better explains the behavioral data compared to the RW model (BIC_RW_=97.6±5.9, BIC_RW±_=89.8±6.0), even accounting for its additional degree of freedom (t(34)= 2.6414, p=0.0124, paired t-test (**Table 1** and **Fig. S2**).

To confirm that the asymmetry of learning rates is not a particularity of our first experiment, in which the worst possible outcome (“bad news”) was represented by a reward omission, we performed a two-way ANOVA with Experiment (1 and 2) as between subject factor and learning rate type (α^+^ and α^-^) as within-subject factor. The analysis showed no significant effect of experiment (F(1,83)=0.077, P=0.782) and no significant valence x experiment interaction (F(1,83)=3.01, P=0.0864) indicating that the two experiments were comparable, and, if anything, the effect size was bigger in presence of punishments. We found indeed a significant main effect of valence (F(1,83)=29.03, P<0.001) on learning rates. Accordingly, post-hoc test revealed that α^-^ was significantly smaller than α^+^ also in the second experiment (t(34)=3.8639, p<0.001 paired t-test) (**Fig. S3A**). These results confirm that optimistic reinforcement learning is not particular to situations involving only rewards but it is still maintained in situations involving both rewards and punishments.

## Discussion

We found that, in a simple instrumental learning task involving neutral visual stimuli associated to actual monetary rewards, participants preferentially updated option values following better-than-expected, compared to worse-than-expected, outcomes. This learning asymmetry was replicated in two experiments and proved to be robust across different conditions.

Our results support the hypothesis that the good news/bad news stands as a core psychological process generating and maintaining unrealistic optimism^8^. In addition, our study has the originality of showing that this effect is not specific to probabilistic belief updating, and that the good news/bad news effect can parsimoniously be considered as an amplification of a primary instrumental learning asymmetry. In other terms, following nomenclature recently proposed by Sharot and Garrett, we found that asymmetric update applies to “prediction errors” and not only to “estimation errors”, as reported in previous studies^9^. Recently, an animated debate emerged concerning whether or not the good news/bad news effect is an artifact due to the fact that in the original task the prospects were very rare life events^22,23^. Our results, by showing that the learning asymmetry persists for abstract cues (as opposite to rare events) associated with not extremely low (nor extremely high) reward probabilities, significantly adds to this debate.

The asymmetric model (RW±) included two different learning rates following positive and negative prediction errors and we found the “positive” learning rate higher compared to the “negative” one^24,25^. A point, which is worth noting, is that optimism seems not to come from overemphasizing gains, but underestimating losses. Note the fact that the learning asymmetry was replicated when the negative prediction errors (i.e. “bad news) were associated with both reward omissions (Experiment 1) and monetary punishments (Experiment 2) indicating that our results cannot be interpreted as a consequence of different processing of outcome values^26^. In other terms the learning asymmetry is not outcome sign-based, but prediction error sign-based.

In principle RW± subjects could have displayed both an optimistic and a pessimistic update, meaning that the ∆BIC is not – *a priori* – a measure of optimism. However, in the light of our results, this metric was *a posteriori* associated with the good news/bad news effect at the individual level. Categorizing subjects based on the ∆BIC, instead of the learning rate difference, has the advantage that the learning rate difference can take positive and negative values in RW subjects, but this difference merely only captures noise, because it is not justified by model comparison. Our subject categorization was further supported by unsupervised Gaussian-mixtures analysis, which indicated that 1) two clusters better explained the data compared to one cluster and that 2) the two cluster corresponded to positive and negative ∆BIC respectively. The combination of individual model comparison with clustering techniques may represent a useful practice for computational phenotyping and for investigating inter-individual cognitive differences^27^.

A higher learning rate for positive compared to negative prediction errors was not the only computational metric distinguishing optimistic from unbiased subjects. In fact, we also found that optimistic subjects had a greater tendency to exploit previously rewarded option, as opposed to unbiased subjects who were more prone to explore both options. Importantly the higher stochasticity of unbiased subjects was associated neither with lower performance in the asymmetrical conditions, nor with a lower baseline quality of fit, as measured by the maximum likelihood. This overexploitation tendency was particularly striking in the symmetrical 25/25% condition, in which both options are poorly rewarding compared to the average task reward rate.

Whereas some previous studies suggest that optimists are more likely to explore and take risks (i.e. entrepreneurs)^28^, we found an association between optimistic learning and higher propensity to exploit. Indeed, the tendency to ignore negative feedback about chosen options was linked to considering a previously rewarded option better than it is, and hence to stick to this preference. A possible link between optimism and such “conservatism” is not new; it can be dated back to Voltaire’s work “*Candide ou l’Optimisme*”, where the belief of “living in the best of the possible worlds” was consistently associated with a strong rejection and condemnation of progress and explorative behavior. In the words of the 18^th^ century philosopher:

*"Optimism," said Cacambo, "What is that?" "Alas!' replied Candide, "It is the obstinacy of maintaining that everything is best when it is worst"^†^*

Accordingly, optimism bias has been recently recognized as an important psychological factor helping maintain inaction regarding pressing social problems, such as climate changes^29^.

Recent studies investigated the neural implementation of the good news/bad news effect when analyzed in the context of probabilistic belief updating. At the functional level, decreased belief updating after worse-than-expected information has been associated with a reduced neural activity in the right inferior prefrontal gyrus (IFG)^7^. Subsequent studies from the same group also showed that boosting dopaminergic function increases the good news/bad news effect and that this bias is correlated with striatal white matter connectivity, suggesting a possible role for the brain reward system^13,30^. Accordingly a more recent study showed differences in the reward system, including the striatum and the ventromedial prefrontal cortex^31^. Consistent with these results, we found that reward prediction error encoded in the brain reward network, including the striatum (mostly its ventral parts) and the vmPFC, was higher in optimistic, compared to unbiased, subjects. Replicating previous findings, we also found a neural network, encompassing the dmPFC and the anterior Insula negatively representing chosen option value^32,33^. When comparing between the two groups we found no difference between optimists and pessimists in this decision-related area ^34,35^. Our results suggest that at the neural level outcome-related activity discriminates between optimistic and unbiased subjects. Remarkably, by identifying functional differences between the two groups, our imaging data corroborates our model comparison-based classification of subject (computational phenotyping).

An important question is unanswered by our study and remains to be addressed. Whereas our results clearly show an asymmetry in the learning process, we cannot decide whether the learning process itself involves the representational space of values or that of probabilities. This question is related to the broader debate whether the reinforcement or the Bayesian learning framework better captures learning and decision-making: two views that have been hard to disentangle, because of largely overlapping predictions, both at the behavioral and neural levels^36–38^. Our results cannot decide whether this optimistic bias is a valuation or a confirmation bias. In other terms, do subjects preferentially learn from positive outcome because of its valence or because a positive outcome “confirms” the choice subjects just made? Future studies, decoupling valence from choice, are required to disentangle these two hypotheses.

It is worth noting that whereas some previous studies reported similar findings^39,40^, another one reported the opposite pattern^41^. The difference between the aforementioned study and ours might rely on the fact that the former involved Pavlovian conditioning. It may therefore be argued that optimistic reinforcement learning is specific to instrumental (as opposite to classical) conditioning.

A legitimate question is why such learning bias survived in the course of evolution? An obvious answer to this question is that being (unrealistically) optimistic is and/or has been, at least in certain conditions, adaptive, meaning that it confers an advantage. Consistent with this idea, in everyday life dispositional optimism^42^ has been linked for instance to better global emotional well-being, interpersonal relationship or physical health. Optimists are less likely to develop coronary heart disease^43^, have broader social network^44^ and are less subject to distress when facing adversity^42^. Over-confidence in one own performance has been shown to be associated with higher performances in competitive games^45^. Such advantages of dispositional optimism could explain, at least in part, the pervasiveness of an optimistic bias in human. Concerning the specific context of optimistic reinforcement learning a recent paper^46^ by Cazé and al. showed that in certain conditions (low rewarding environments), an agent learning asymmetrically in an optimistic manner (i.e. with a higher learning rate for positive than for negative feedback) objectively outperforms another “unbiased” agent in a simple probabilistic learning task. Thus, before any social, well-being or health consideration, it is normatively advantageous (in certain contingencies) to take more into account positive than negative feedback. Thus a possible explanation for an asymmetric learning system is that the conditions identified by Cazé et al. closely resemble to the statistics of the natural environment that shaped the evolution of our learning system.

Finally, when reasoning about the adaptive value of optimism, a crucial point to take into account is the significant inter-individual variability of unrealistic optimism^7,11–13^. As social animals, humans face both private and collective decision-making problems^47^. An intriguing possibility is that multiple “sub-optimal” reinforcement learning strategies are maintained in the natural population to ensure an “optimal” learning repertoire, flexible enough to solve at the group-level the value learning and exploration-exploitation tradeoff^48^. This hypothesis needs to be formally addressed using evolutionary simulations.

To conclude, our findings shed new light on the nature of the good news/bad news effect and therefore on the mechanistic origins of unrealistic optimism. We suggest that the optimistic learning is not specific to “high-level” belief updating but a particular consequence of a more general “low-level” instrumental learning asymmetry, which is associated to enhanced prediction error encoding in the brain reward system.

### Materials and Methods

#### Subjects

The first dataset (N=50) served as a cohort of healthy control subjects in a previous clinical neuroimaging study^15^. The second dataset involved the recruitment of new subjects (N=35). The local ethics committees approved both experiments. All subjects gave written informed consent before inclusion in the study and the study was carried out in accordance with the declaration of Helsinki (1964, revised 2013). In both studies the inclusion criteria were being older than 18 years and having no history of neurologic or psychiatric disorders. In experiments 1 and 2, men / women ratios were 27/23 and 20/15 respectively and the age means 27.1 ± 1.3 and 23.5 ± 0.7 respectively (expressed as mean ± S.E.M). In the first experiment subjects believed that they would be playing for real money, the final payoff was rounded up to a fixed amount of 80€ for every participant. In the second experiment subjects were paid the exact amount of money earned in the learning task, plus a fixed amount (average payoff 15.7±7.6€).

#### Behavioral task and analyses

Subjects performed a probabilistic instrumental learning task described previously^16^ (**Fig. 1A**). Briefly, the task involved choosing between two cues that were associated with stationary reward probability (25% or 75%). There were 4 pairs of cues, randomly constituted and assigned to the 4 possible combinations of probabilities (25/25%, 25/75%, 75/25%, and 75/75%). Each pair of cues was presented 24 times, each trial lasted in average 7000ms. Subjects were encouraged to accumulate as much money as possible and were informed that some cues would result in a win more often than others (the instructions have been published in appendix of the original study^16^). Subjects were given no explicit information regarding reward probabilities, which they had to learn through trial and error. The positive outcome reward was winning money (+0.50€); the negative outcome was getting nothing (0.0€) in the first experiment and losing money (-0.50€) in the second experiment. Subjects made their choice by pressing left or right response buttons with a left or right hand finger. Two given cues were always presented together, thus forming a fixed pair (choice context).

Regarding payoff, learning mattered only for pairs with unequal probabilities (75/25% and 25/75%). As dependent variable we extracted the correct response rate in asymmetric conditions (i.e. the left response rate for the 75/25% pair and the right response rate in the 25/75% pair) (**Fig. 1B**). In symmetrical reward probability conditions, we calculated the so-called “preferred response rate”. The preferred response was defined as the most chosen option, i.e. chosen by the subject more than 50% of the trials. This quantity is therefore, by definition greater than 50%. The analyses focused on the preferred choice rate in the low reward condition (25/25%), where standard models predict greater frequency of negative prediction errors. Behavioral variables were compared within-subjects using paired two-tailed t-test and between-subjects using two-sample two-tailed t-test. Interactions were assessed using ANOVA.

#### Computational models

We fitted the data with reinforcement learning models. The model space included a standard Rescorla-Wagner model (or Q-learning)^18,19^ (thereafter referred to as RW) and a modified version of the latter accounting differentially for learning from positive and negative prediction errors (thereafter referred to as RW±)^25,40^. For each pair of cues, the model estimates the expected values of left and right options, *Q*_*L*_ and *Q*_*R*_, on the basis of individual sequences of choices and outcomes. These Q-values essentially represent the expected reward obtained by taking a particular option in a given context. In the first experiment, that involved only reward and reward omission, Q-values were set at 0.25€ before learning, corresponding to the a priori expectation of 50% chance of winning 0.5€ plus a 50% chance of getting nothing. In the second experiment, which involved reward and punishment, Q-values were set at 0.0€ before learning, corresponding to the a priori expectation of 50% chance of winning 0.5€ plus 50% chance of losing 0.5€. These priors on the initial Q-values are based on the fact that first subjects were explicitly told in the instruction that no symbol was deterministically associated to either of the two possible outcomes and on the fact that subjects were implicitly exposed to the average task outcome during the training session. Further control analyses using post training (“empirical”) initial Q-values, have been performed and are presented in the **Supplementary Materials** and **Fig. S6**. After every trial *t*, the value of the chosen option (e.g., L) was updated according to the following rule:

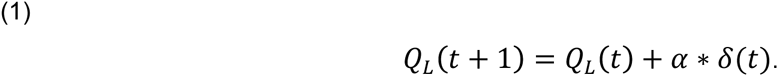

In the equation, δ(*t*) was the prediction error, calculated as:

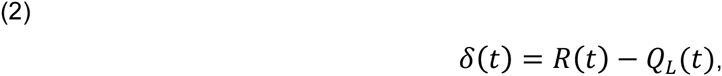

and *R*(*t*) was the reward obtained as an outcome of choosing *L* at trial *t*. In other words, the prediction error δ(*t*) is the difference between the expected reward *Q*_*L*_(*t*) and the actual reward *R*(*t*). The reward magnitude *R* was +0.5 for winning 0.5€, 0 for getting nothing and -0.5 for losing 0.5€. The learning rate, *α*, is a scaling parameter that adjusts the amplitude of value changes from one trial to the next. Following this rule, option values are increased if the outcome is better than expected and decreased in the opposite case and the amplitude of the update is similar following positive and negative prediction errors.

The modified version of Q-Learning algorithm (RW±) differs from the original one (RW) by its *Q* values updating rule, as follows:

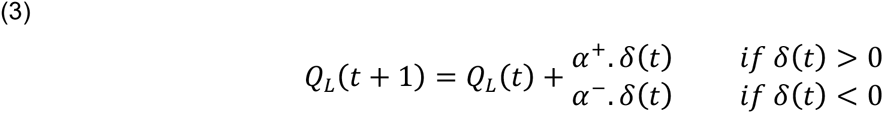

The learning rate *α*^+^ adjusts the amplitude of value changes from one trial to the next when prediction error is positive (when the actual reward *R*(*t*) is better than the expected reward *Q*_*L*_(*t*)) and the second learning rate *α*^-^ does the same when prediction error is negative. Thus the RW± model allows for the amplitude of the update being different, following positive (“good news”) and negative (“bad news”) prediction errors and permits to account for individual differences in the way subjects learn from positive and negative experience. If both learning rates are equivalent, *α*^+^ = *α*^-^, RW± model equals the RW model. If *α*^+^ > *α*^-^, subjects learn more from positive than negative events. We refer to this case here as optimistic reinforcement learning. If *α*^+^ < *α*^-^, subjects learn more from negative than positive events. We refer to this case here as pessimistic reinforcement learning (**Fig. 2B**).

Finally, given the Q-values, the associated probability (or likelihood) of selecting each option was estimated by implementing the soft-max rule for choosing *L*, which is as follows:

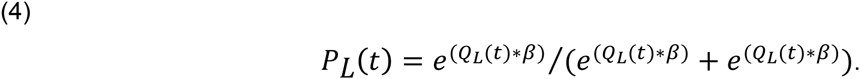

This is a standard stochastic decision rule that calculates the probability of selecting one of a set of options according to their associated values. The temperature, *β*, is another scaling parameter that adjusts the stochasticity of decision-making and by doing so controls the exploration/exploitation trade-off.

#### Model comparison

We optimized model parameters by minimizing the negative log-likelihood of the data given different parameters settings using Matlab’s fmincon function, as previously described^49^. Additional parameter recovery analyses based on model simulations show that our parameter optimization procedure correctly retrieves parameters’ values (**Supplementary Materials** and **Fig. S7**). Negative log-likelihoods (LLmax) were used to compute at the individual level (random effects) for each model the Bayesian information criterion as follows:

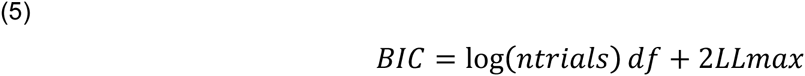

We computed then the inter-individual average BIC in order to compare the quality of the fit of the two models, while accounting for their difference in complexity. The intra-individual difference in BIC (∆BIC=BIC_RW_ - BIC_RW±_) was also computed in order to categorize subjects in two groups and to quantitatively describe at the individual level the divergence from an unbiased model (**Fig. 2A**): RW± subjects, whose ∆BIC is positive, are better explained by RW± model. RW subjects, whose ∆BIC is negative, are better explained by RW model. We note that lower BIC indicated better fit. We also calculated the model exceedance probability and the model expected frequency based on the BIC as an approximation of the model evidence. (**Table 1**). Individual BIC were fed into the mbb-vb-toolbox, a procedure that estimates the expected frequencies and the exceedance probability for each model within a set of models, given the data gathered from all participants. Exceedance probability (denoted XP) is the probability that a given model fits the data better than all other models in the set.

The model parameters, (*α*^+^, *α*^-^ and 1/*β*) were also compared between the two groups of subjects. Learning rates were compared using a mixed ANOVA with group (RW vs RW±) as a between-subject factor and learning rate type (+ or -) as a within-subject factor. The temperature was compared using a two-sample two-tailed t-test. The normalized learning rates asymmetry (α^+^ - α^-^) / (α^+^ + α^-^) was also computed as a measure of the good news/bad news effect and used to assess correlation with behavioral and neural data.

### Subject classification

Subjects were classified based on the ∆BIC, which is the intra-individual difference in BIC between the RW and RW± model. While controlling for model parsimony, positive value indicates that the RW± better fits the data; negative value indicates the RW model better fit. The cut-off of ∆BIC=0 is *a priori* meaningful because it indicates the limit beyond which there is enough (Bayesian) evidence to consider that a given subject’s behavior corresponds to a more complex model involving two learning rates. We also validated the <u>∆BIC=</u>0 cut-off *a posteriori* with unsupervised clustering. We fitted Gaussian mixed distributions to individual ∆BICs (N=85, corresponding to the two experiments) using MatLab function gmdistribution.m. The analysis indicated that two clusters significantly better explain the variance compared to one cluster (k=1, BIC = 716.4; k=2 BIC = 635.6). The two clusters largely corresponded to subjects with negative (N=40, min= -6.4; mean= -3.6, max= -0.9) and positive ∆BIC (N=45, min=-0.5, mean=15.7, max=60.6). The two cluster differed in both the normalized difference in learning rates (0.14 vs. 0.73; t(83)=7.2, P<0.001) and exploration rate (0.32 vs 0.09; t(83)=7.2, P=0.006).

### Model simulations

We also analyzed the models’ generative performance by the mean of model simulations. For each participant we devised a virtual subject, represented by a set of individual best fitting parameters. Each virtual subject dataset was obtained averaging 100 simulations, to avoid any local effect of the individual history of choice and outcome. The model simulations included all task conditions. The evaluation of generative performances involved the assessment of the “winning model’s” ability to reproduce the key statistical effects of the data, as opposite to the “losing model”. Unlike Bayesian model comparison, model simulation comparison is bounded to a particular behavioral effect of interest (in our case the preferred response rate). The model simulation analysis, which is focused on the evidence “against” a given model, is complementary to the Bayesian model comparison analysis, which is focused on the evidence in favor of a model^50,51^.

### Imaging data Acquisition & Analysis

Subject of the first experiment (N=50) performed the task magnetic resonance imaging (MRI) scanning. T1-weighted structural images and T2*-weighted echo planar images (EPIs) were acquired during the first experiment and analyzed with the Statistical Parametric Mapping software (SPM8; Wellcome Department of Imaging Neuroscience, London, England). Acquisition and preprocessing parameters were previously and extensively described^15,16^. We refer to these publications for details about image acquisition and preprocessing.

### Functional magnetic resonance imaging analysis

The fMRI analysis was based on a single general linear model. Each trial was modeled as having two time points, stimuli and outcome onsets. Each time point was regressed with a parameter modulator. Stimuli onset was modulated by the chosen option value (*Q*_*Chosen*_(*t*)); outcome onset was modulated by the reward prediction error δ(*t*)). Given that different subjects did not implement the same model, the choice of the model used to generate the parametric regressors is not obvious. Since the RW± and the RW models are nested and the RW± model was the group-level best fitting model, we opted for using its parameters to generate the regressors. However, note that confirmatory analyses using for each group its best fitting model’s parameters lead to similar results. The parametric modulators were z-scored to ensure between subject scaling of regression coefficients^52^. Linear contrasts of regression coefficients were computed at the subject level and compared against zero (one-sample t-test). Statistical parametric maps were threshold at p<0.05 with a voxel-level family-wise error (FWE) correction and a minimum of 60 contiguous voxels. Whole brain analysis was performed including both group of subjects and leads to the identification of functionally characterized neural networks used to define unbiased ROIs. The dmPFC and the Insular ROIs were defined as the intersection of the voxels significantly correlating with Q_Chosen_(t) and aal (i.e. automatic anatomical labeling) masks of the medial frontal cortex (including the superior frontal gyrus, the SMA and the anterior medial cingulate) and the bilateral insula, respectively. The vmPFC and the striatal ROIs were defined as the intersection of the voxels significantly correlating with δ(t) and aal (i.e. automatic anatomical labeling) masks of the ventral prefrontal cortex (including the anterior cingulate, the gyrus rectus and the superior frontal gyrus, orbital part and medial orbital part) and the bilateral caudate and putamen, respectively. Within ROIs the regression coefficients were compared between-group using two-sample two-tailed t-test.

**Fig. 4:**
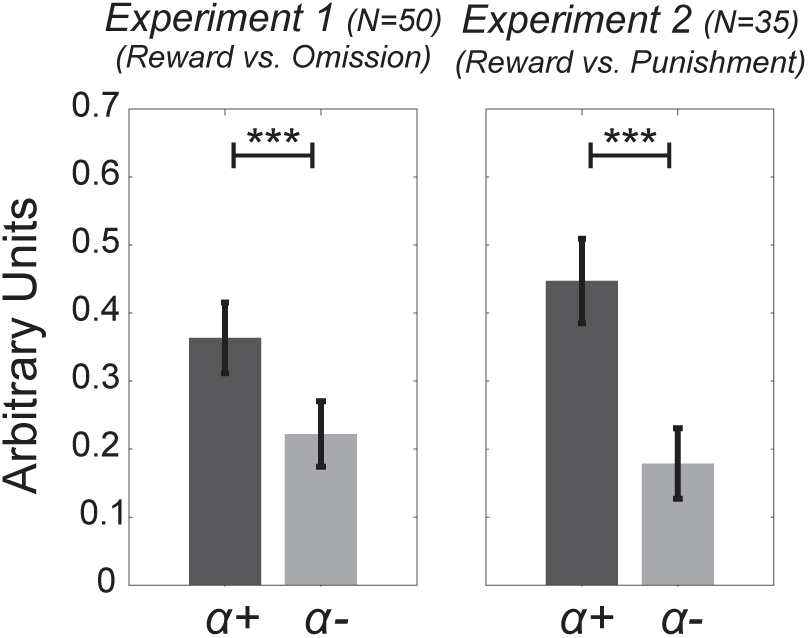
Robustness of optimistic reinforcement learning. Histograms show the learning rates following positive prediction errors (α^+^) and negative prediction errors (α^-^), in Experiment 1 (N=50) and Experiment 2 (N=35). Experiment 1’s worst outcome was getting nothing (0€). Experiment 2’s worst outcome was losing money (-0.50€).

## Acknowledgments

The authors acknowledge Yulia Worbe and Mathias Pessiglione for granting access to the first dataset. We thank Valentin Wyart, Bahador Bahrami, Bojana Kuzmanovic and the anonymous reviewers for their helpful comments. We thank Tali Sharot and Neil Garrett for kindly providing activation masks. SP was supported by a Marie Sklodowska-Curie Individual European Fellowship (PIEF-GA-2012 Grant 328822) and is currently supported by an ATIP-Avenir grant. GL was supported by a PHD fellowship of the Ministère de l'enseignement supérieur et de la recherche. ML was supported by an EU Marie Sklodowska-Curie Individual Fellowship (IF-2015 Grant 657904) and acknowledges the support of the Bettencourt-Schueller Foundation. The second experiment was supported by the ANR-ORA, Nesshi 2010-2015 research project to SBG.

## Footnotes

* Bacon, F. (1939). *Novum organum*. In Burtt, E. A. (Ed.), The English philosophers from Bacon to Mill (pp. 24-123). New York: Random House. (Original work published in 1620)

† Original French citation: "*Qu’est-ce qu’optimisme? disait Cacambo. – Hélas! dit Candide, c’est la rage de soutenir que tout est bien quand on est mal*." Voltaire (2014), *Candide ou l'optimisme*, Arvensa editions, p56, Ch. XIX. (Original work published in 1759)

## Supplementary Information

### Preferred response rate as a behavioral measure of optimistic behavior

As previously defined in the main text, the preferred response rate is the rate of the choices directed toward the most frequently chosen option by subjects in symmetric reward probability conditions (i.e. 25/25% and 75/75%). The preferred choice rate is therefore by definition greater than 0.5. In these conditions there is no contingency-based reason to prefer one option to the other. This is particularly true in low-rewarding environment (25/25% condition), where neither option is satisfying in terms of outcome, compared to the average task outcome. We showed previously that the preferred response rate allows to behaviorally differentiating optimistic from unbiased subjects.

RW± subjects were characterized at the computational level by two features that are good news/bad news effect (i.e. learning rate asymmetry) and lower exploration rate. Both these computational features concur to generate this behavioral pattern. **Fig. S1A** shows the preferred choice rate in the 25/25% condition of a typical RW± subject, whose behavior is much better explained by the RW± model (∆BIC=58.2).

**Fig. S1:**
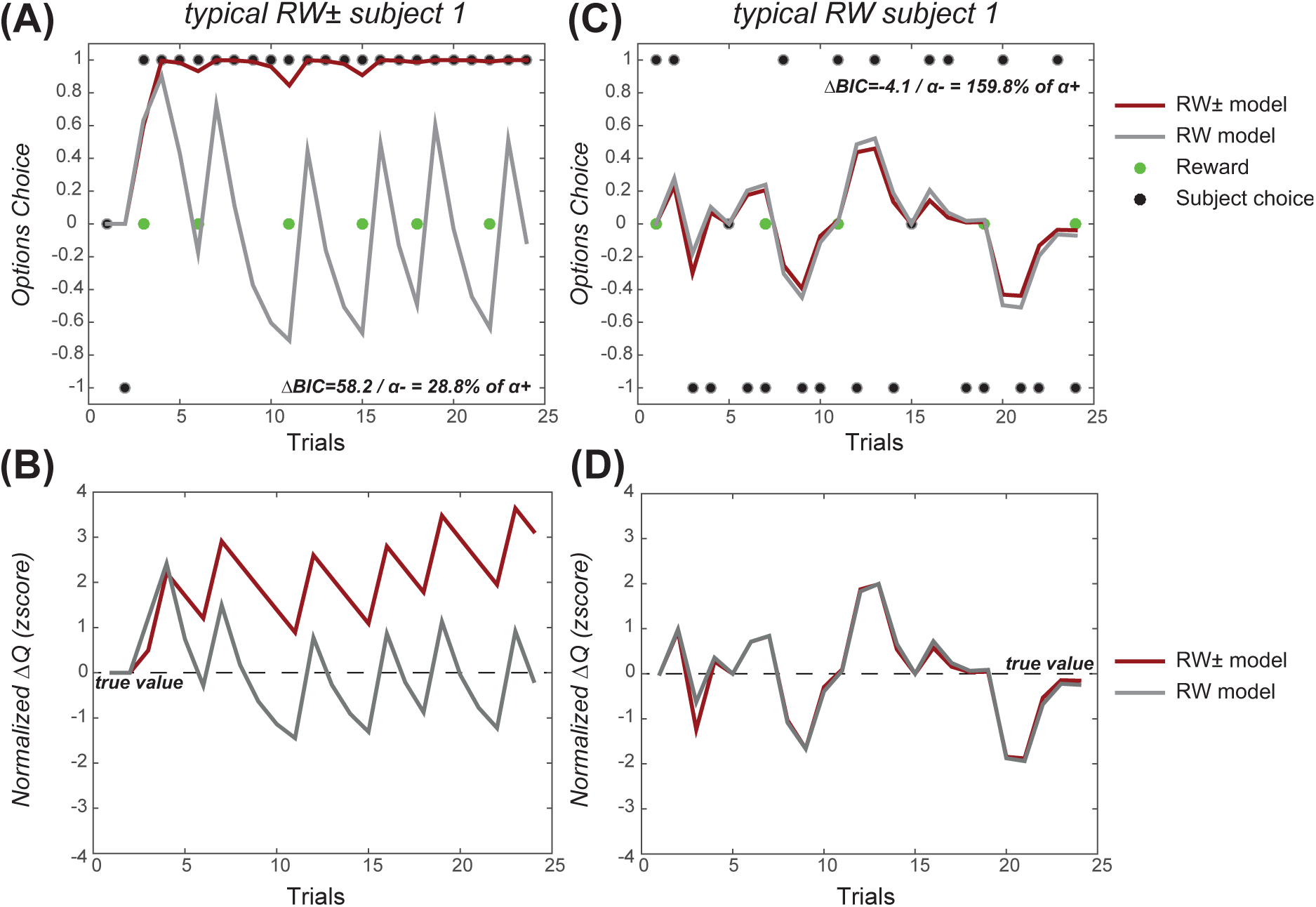
typical “optimistic” and “unbiased” subjects in the 25/25% condition. **(A)** and **(B)** RW± (optimistic) typical subject. (**A**) Plot represents behavioral choices (represented by black dots) of a typical RW± subject (i.e. whose behavior is best fitted by the RW± model) in the 25/25% condition, together with RW and RW± models predictions (represented respectively by gray and red lines). (**B**) Plot represents Q-values (of the two options) differential evolution in each model for a typical RW± subject. **(C)** and **(D)** RW (unbiased) typical subject. **(C)** Plot represents behavioral choices (represented by black dots) of a typical RW subject (whose behavior is best fitted by the RW model) in the 25/25% condition, together with RW and RW± models predictions (represented respectively by gray and red lines). (**D**) Plot represents the evolution of Q-values (of the two options) differential in each model for a typical RW subject.

We clearly see that his choices are stabilized toward one option (preferred response rate=0.94), after one single reward event. RW± model fit captures this preference, by giving less weight to negative feedback than to positive one (α^-^ is approximately four times smaller than α^+^) and by allowing a very little exploration rate (1/β=0.001). This learning rate asymmetry creates and accentuates over the trials the “preferred minus non-preferred” ∆Q (whose true value is zero), which is further reinforced by not exploring the other option (**Fig.S1B**). At the opposite (**Fig.S1C**), a typical unbiased subject (RW) does not show clear preference toward one of the options (preferred response rate around 0.58). Accordingly the RW model better explains his behavior (∆BIC=-4.1) and his learning rate asymmetry is moderate (α^-^ is approximately 1.6 time higher than α^+^ and the exploration rate higher (1/β=0.235). This symmetry between positive and negative learning rates and the tendency to extensively explore the two options do not give advantage to any of the Q-Values, the differential of which gravitates around zero over learning (i.e its true value) (**Fig.S1B**). These examples nicely illustrate how the preferred response rate is affected by learning rate asymmetry and exploration rate thus allowing discriminating RW± and RW subjects.

In order to further illustrate how the preferred response rate relates to both computational signatures of optimistic reinforcement learning, we run model simulations distinguishing the effect of the latter two features, a positive learning rates asymmetry and a low exploration rate. We ran four simulations by experiment using parameters from either typical RW subjects or typical RW± subjects. The simulations were generated using the parameter sets of both experiments. In the simulations based on Experiment 1 (N=100 virtual subjects) we used either symmetric learning rates (RW: α^+^ = α^-^= 0.41) or optimistically asymmetric (RW± : α^+^ = 0.27 and α^-^ = 0.04) and the exploration rate was either low (RW±: 1/β = 0.06) or high (RW: 1/β = 0.21). In the simulations based on Experiment 2 (N=105 virtual subjects) we used either symmetric learning rates (RW: α^+^ = α^-^ = 0.27) or optimistically asymmetric (RW±: α^+^ = 0.47 and α^-^ = 0.10) and the exploration rate was either low (1/β = 0.15) or high (1/β = 0.73). Results from those simulations presented in the **Fig. S2**, showed that neither of the two computational features alone (learning rate asymmetry and lower exploration rate) are sufficient to reach the preferred response rate of RW± subjects. On the contrary, simulations ran with both computational features permit to the preferred response rate to be very close to the empirical results of RW± subjects. In other terms, the learning rate asymmetry and the exploration rate have as super-additive effect on the preferred response rate, which is an essential characteristic of the RW± (optimistic) phenotype.

**Fig. S2:**
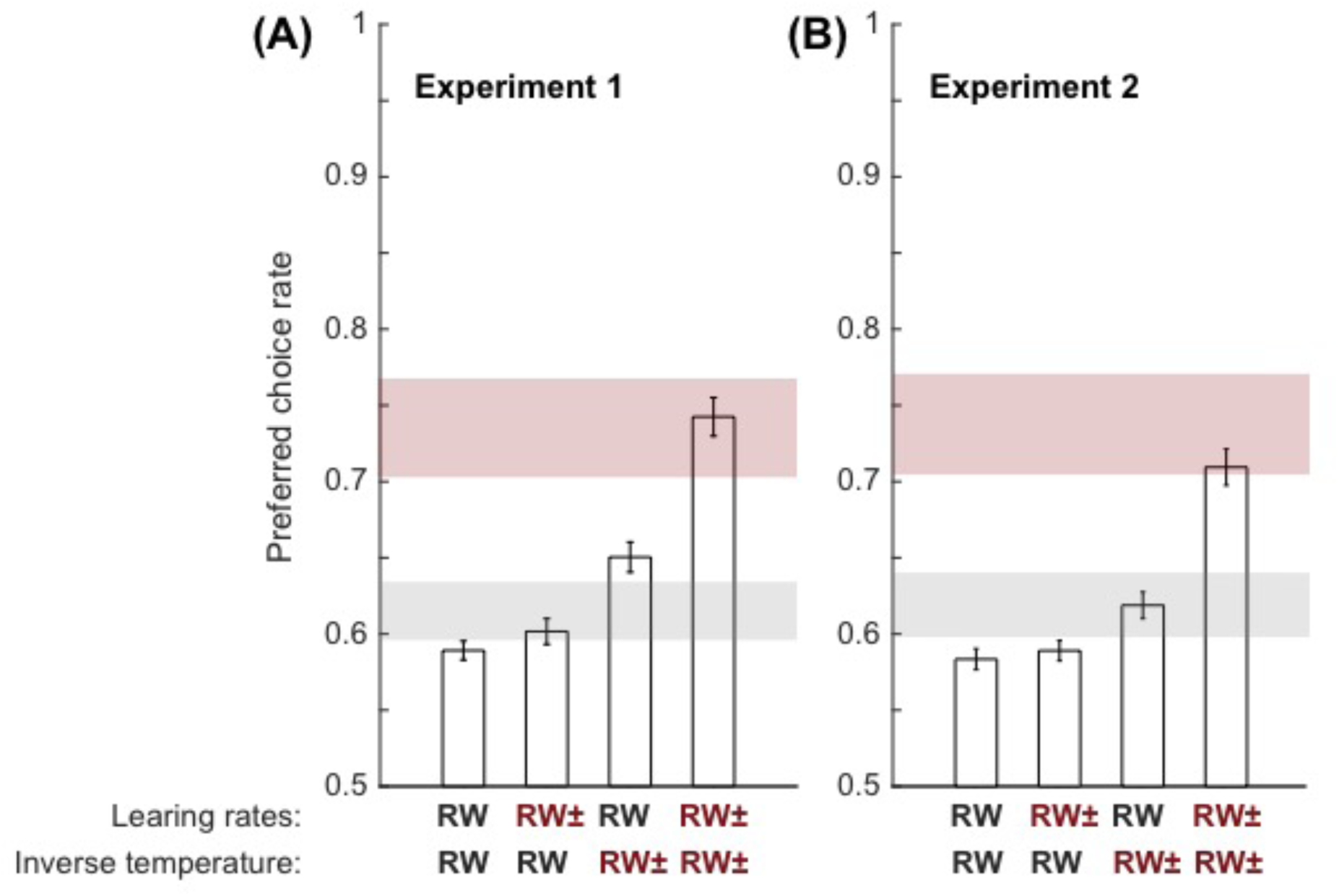
contribution of the learning rate asymmetry and the choice inverse temperature to the preferred choice rate. **(A)** and **(B)** Bars represent the simulated preferred response rate in four conditions varying according to the model parameters used to simulate the data. RW learning rates are symmetrical and correspond to the average learning rates of RW subjects while RW± learning rates are asymmetrical and correspond to the average learning rates of RW± subjects. The RW inverse temperature is high and matches the average inverse temperature of RW subjects while RW± inverse temperature is low and matches the average inverse temperature of RW± subjects. Finally, the horizontal colored areas represent the average empirical preferred response rate plus or minus its standard deviation to the mean, in RW subjects (grey area) and RW± subjects (red area).

### Optimistic reinforcement learning is robust across different learning phases

Previous studies have shown that learning rates adapt with learning. More precisely the learning rate may be reduced when the confidence about choice’s outcome is high and, conversely, augmented in situation of high uncertainty^1,2^. It could be argued that in our task the optimistic learning rate asymmetry (α^+^>α^-^ was specifically driven by the late trials, when the reward contingences have been learnt and the subject has no longer need to monitor prediction errors as objectively as in the early trials. In order to assess that the learning rate asymmetry was not specific of the late learning, we analyzed and compared RW± model parameters separately optimized in the first and second halves of the task in both experiments (**Fig. S3A**). We performed a two-way ANOVA with part of the task (first and second halves) and learning rate type (α^+^ and α^-^) as within-subject factors. It shows no significant effect of task period (F(1,84)=0.011, P=0.917) and no significant valence x period interaction (F(1,84)=0.011, P=0.917) indicating that the learning rates asymmetry is not specific to the late trials. We found indeed a significant main effect of valence (F(1,84)= 46.42, P<0.001) on learning rates. Accordingly, post-hoc test revealed that α^-^ was significantly smaller than α^+^ also in the first half (t(84)=5.7214, p<0.001 paired t-test) and in the second half (t(84)= 5.3764, p<0.001 paired t-test) of the task.

**Fig. S3:**
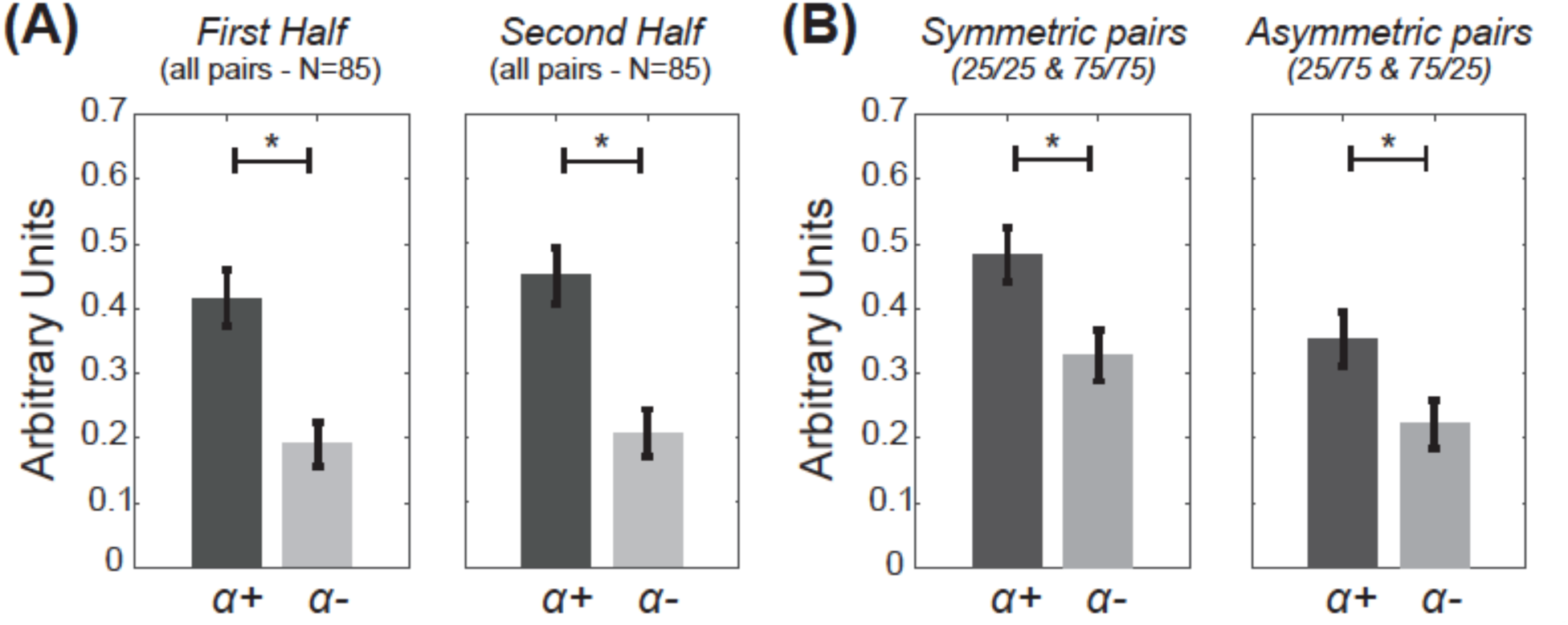
Robustness of optimistic reinforcement learning. **(A)** Control first half / second half. Histograms show the learning rates following positive prediction errors (*α*^+^) and negative prediction errors (*α*^-^), obtained from parameters optimization separately performed in the first and second halves of the experiments (N=85). **(B)** Control symmetric / asymmetric conditions. Histograms show the learning rates following positive prediction errors (*α*^-^) and negative prediction errors (*α*^+^), obtained from parameters optimization involving only the “symmetric or the “asymmetric” conditions (N=85).

### Optimistic reinforcement learning is robust across different outcome contingencies

It has also been proposed that learning rates may adapt as a function of task contingencies^3^. In our task the macroscopic (aggregate) model-free signature of optimistic behavior was found in the symmetrical conditions: higher preferred choice rate in the RW± subjects (**Fig. 2** and **S4**). In the main text we reported the results concerning the 25/25% condition, but this was also true for the 75/75% condition, where the preferred choice rate in the RW± subjects was higher compared to the RW subjects (t(83)=3.6686, p<0.001, two-sample t-test) and compared to what was predicted by the RW model (t(42)=16.0292, p<0.001, paired t-test). It might be argued that the learning rate asymmetry we observed was driven by an adaptation of the learning rates specific to the symmetrical conditions, in which there is no true correct response.

In order to verify that the asymmetry of the learning rate was not only expressed in the symmetric conditions (when options are equally rewarding), we optimized learning rates in symmetric and asymmetric conditions independently (**Fig. S3B**). A two-way ANOVA devised with condition type (symmetric and asymmetric) and learning rates valence as within subjects factors, showed a main effect of valence (F(1,84)=21.14, P<0.001) that is consistent with α^+^ being higher compared to α^-^. It also showed a lower effect of condition type (F(1,84)= 9.493, P=0.003) both learning rates being lower in asymmetric conditions, but importantly no significant interaction between valence and condition type (F(1,84)=0.124, P=0.726). Post hoc tests confirm this learning rates asymmetry (α^+^> α^-^ in both condition types (t(84)=3.1106, p=0.003 in asymmetric conditions and t(84)= 3.139, p=0.002 in symmetric conditions, paired t-tests).

**Fig. S4:**
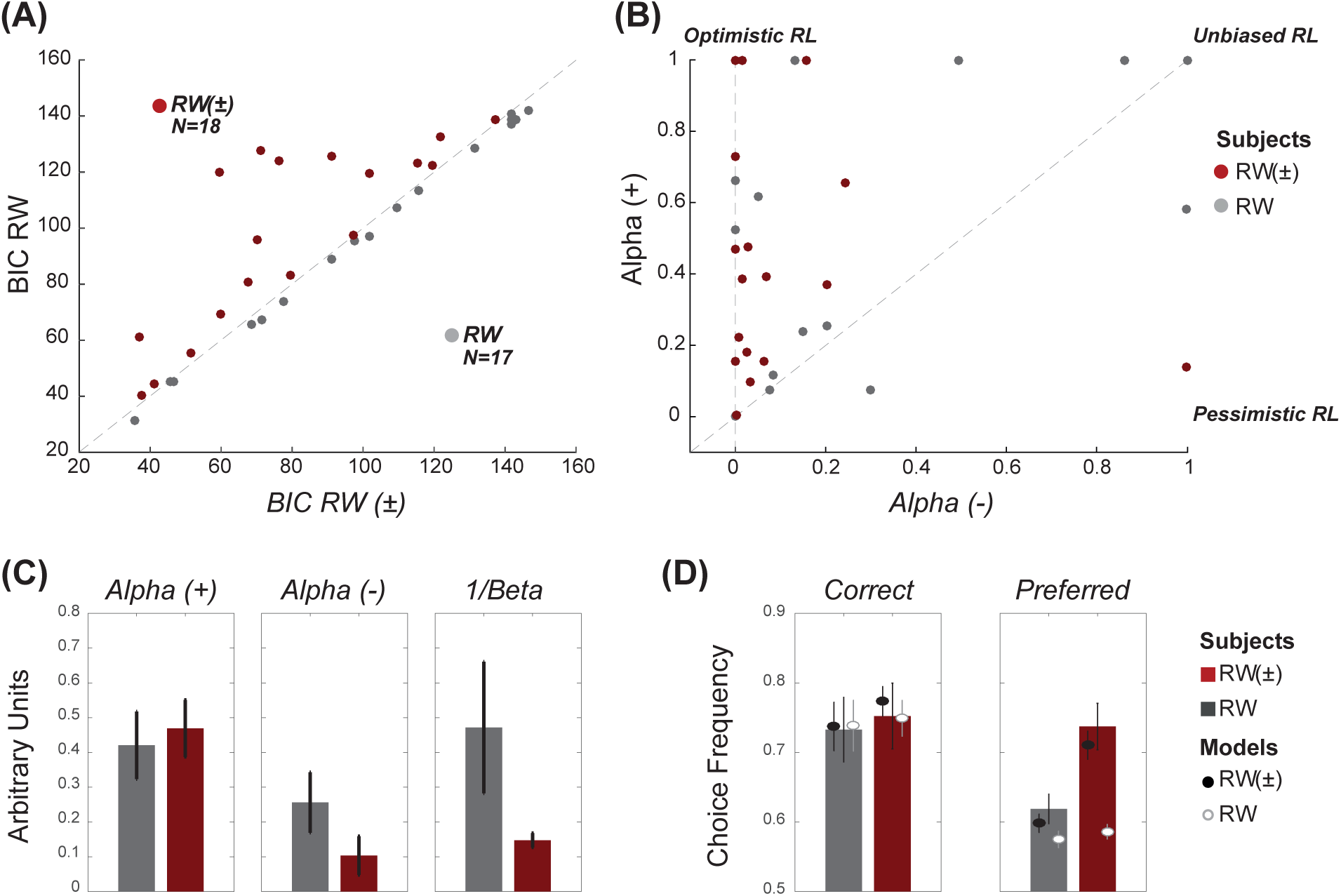
Replication of the computational and behavioral results in another group of subjects and using actual punishments. In order to assess the robustness of optimistic reinforcement learning in presence of actual punishments, we run an additional experiment (Experiment 2; N=35). The probabilistic contingencies, as well as the number of trials, were similar in both experiments. However, whereas Experiment 1’s worst outcome was getting nothing (0€), Experiment 2’s worst outcome was losing money (-0.50€). The two experiments led to strikingly similar behavioral and computational results (**Fig. 2**). **(A)** Model comparison. The graphic displays the scatter plot of the BIC calculated for the RW model as a function of the BIC calculated for the RW± model. Subjects are clustered in two populations according to the BIC difference (∆BIC = BICRW - BICRW±) between the two models. RW± subjects (displayed in red) are characterized by a positive ∆BIC, indicating that the RW± model better explains their behavior. RW subjects (displayed in grey) are characterized by a negative ∆BIC, indicating that the RW model better explains their behavior. **(B)** Model parameters. The graphic displays the scatter plot of the learning rate following positive prediction errors α+ as a function of the learning rate following negative prediction errors α-obtained from the RW± model. “Unbiased” reinforcement learning (RL) is characterized by similar learning rates for both types of prediction errors. “Optimistic” RL is characterized by a bigger learning rate only for positive compared to negative prediction errors. “Pessimistic” RL is characterized by the opposite pattern. **(C)** The histograms represent the RW± model free parameters (the learning rates and the inverse temperature 1/beta) as function of the subjects’ populations. **(D)** Actual and simulated choice rates. Histograms represent the observed and dots represent the model simulated of choices for both populations and both models, respectively for correct option (extracted from asymmetric condition), and from preferred option (extracted from the symmetric condition 25/25%, see **Fig. 1A**).

Indeed, an in depth analysis of correct response rate distribution (**Fig. S5**) showed that both behavioral and simulated response distributions are significantly different between groups although being similar within groups. The analysis focused on the distributions of correct response rates in both asymmetric conditions (25/75% and 75/25%) among real and simulated populations both dichotomized in RW± and RW subjects.

**Fig. S5:**
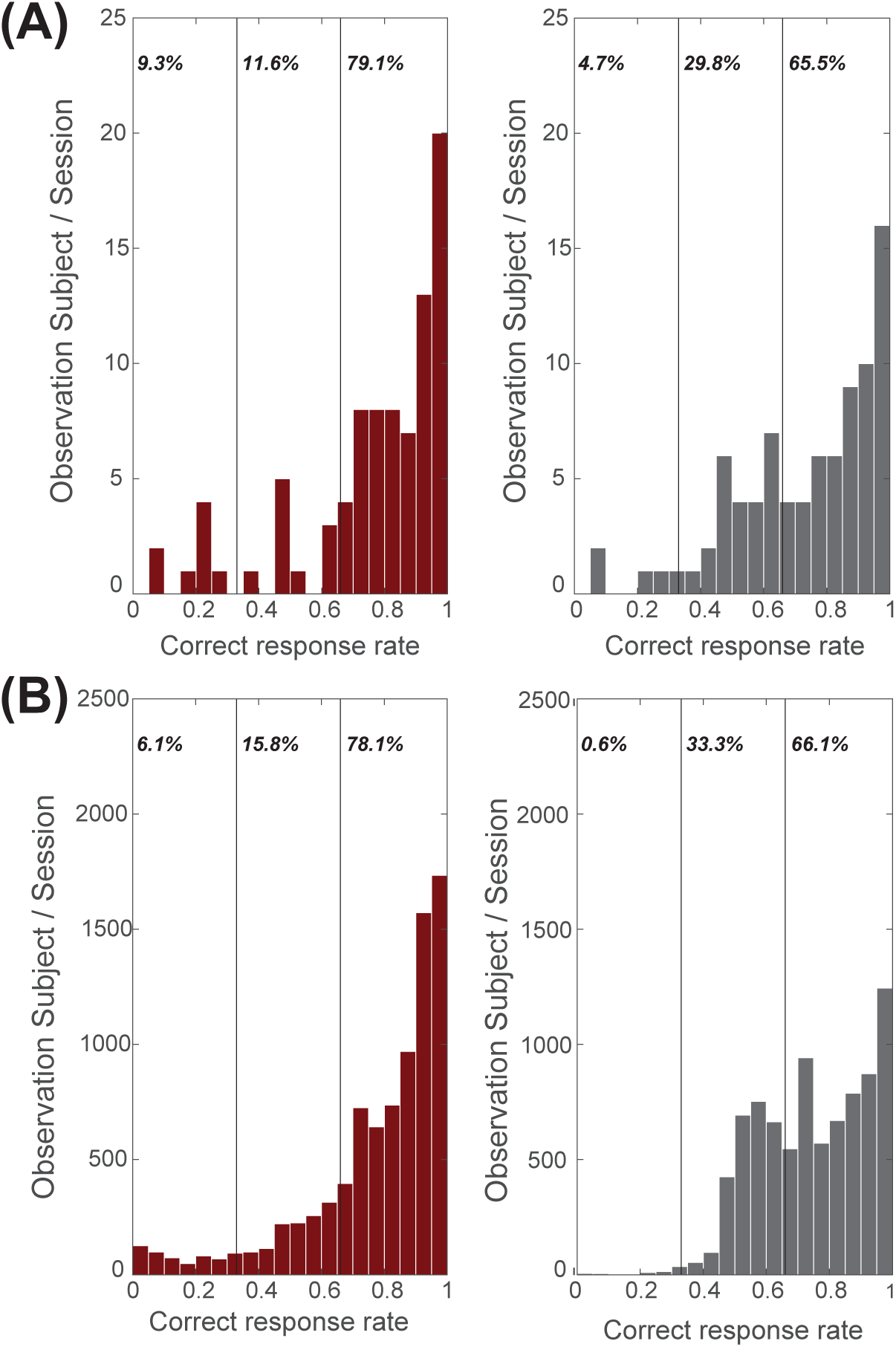
actual and modeled distributions of correct choice frequency. **(A)** Histograms represent distributions of correct choice rate in asymmetric conditions (25/75%, 75/25%) in RW and RW± subjects. Data are taken from both experiments (N=85). **(B)** Histograms represent distributions of correct choice rate in asymmetric conditions (25/75%, 75/25%) in RW and RW± virtual subjects. Each virtual subject (correspond to an individual set of free parameters obtained fitting the actual data with the RW± model) played the task one hundred times (N=8500). RW± real and virtual subjects are characterized by a higher frequency of “extreme” (i.e. greater than 0.66 or lower than 0.33) correct response rate, whereas RW real and virtual subjects, are characterized by a higher frequency of “intermediate” correct response rate.

To compare distributions, we split the correct response rate into three equal categories and calculated the percentage of observations belonging to each category. A first comparison between RW and RW± real populations (**Fig. S5A**) showed that their correct response rate distributions were significantly different (χ^2^=10.69, p<0.005, chi-squared test). The RW± signature here being that distributions are marked by a greater presence of extreme responses due to the sensitivity (greater α^+^ and lower exploration) of RW± subjects to both reward received from the correct option (79.1% and 65.5% respectively), but also to reward accidentally received from the incorrect option (9.3% and 4.7% for RW± and RW respectively). So the insensitivity of RW± subjects to negative feedback and their relatively low tendency to explore available options make them prone to choose and stick with the worst available option. To test whether or not this feature of RW± subjects was captured by the optimistic reinforcement model, we realized the same analysis in a population of virtual RW± et RW subjects (**Fig. S5B**). Firstly, we found that simulated distributions of correct response rate are equivalent to behavioral ones (χ^2^=3.49, p=0.17, for RW group and χ^2^=1.09, p=0.58, for RW± group, chi-squared tests). Secondly and similarly to real distributions, simulated distributions were found to be significantly different between groups (χ^2^=12.66, p<0.005, chi-squared test) with a similar representation of extreme correct response rates in RW± group compared to RW group. Being “optimistic” in this task is then often advantageous for a majority of optimistic subjects displaying a very high rate of correct option. However, when optimists receive a probabilistic reward from the worst option, they can be trapped by their insensitivity to negative feedback and by not being prone to explore alternatives.

### Optimistic reinforcement learning is robust across different Q-values initializations

Learning rates asymmetry was obtained through parameters optimization using original and derivative Q-Learning models. As indicated in the **Methods** section, subjects were induced to have “neutral” priors about each stimulus value via the instructions and the training session. Accordingly, *Q* values were set at 0.25€ before learning, corresponding in the first experiment (that involved only reward +0.5€ and reward omission 0.0€) to the a priori expectation of 50% chance of winning 0.5€ plus a 50% chance of getting nothing. In the second experiment (which involved reward +0.5€ and punishment -0.5€) & *Q* values were set at 0.0€ before learning, corresponding to the a priori expectation of 50% chance of winning 0.5€ plus 50% chance of losing 0.5€. In order to verify the robustness of our result in respect of the Q-value initialization, we performed another parameter optimization using the same models but initializing Q-values using individual “empirical” priors. We defined the “empirical” priors as the average outcome observed during the training session averaged across all the stimuli: we found 0.23±0.004€ (in Experiment 1) and -0.02 ± 0.01€ (in Experiment 2). These values are very close to the theoretical values used in the analysis (0.25€ and 0.00€), except for a small under-estimation that is due to the fact that the worst stimuli are less extensively sampled. Parameters optimized using initial empirical values confirmed, once again, a learning rate asymmetry, consistent with the good news/bad news effect (α^+^ = 0.32±0.06 and α^-^ = 0.17±0.06, t(29)=3.12, p=0.0041 for Experiment 1 and α^+^ = 0.46±0.06 and α^-^ = 0.19±0.05, t(33)=3.73, p<0.001 for Experiment 2) (**Fig S6**). Thus, empirically determined priors were similar to theoretical ones and using the firsts in our parameter optimization procedure have no impact on the results.

**Fig S6:**
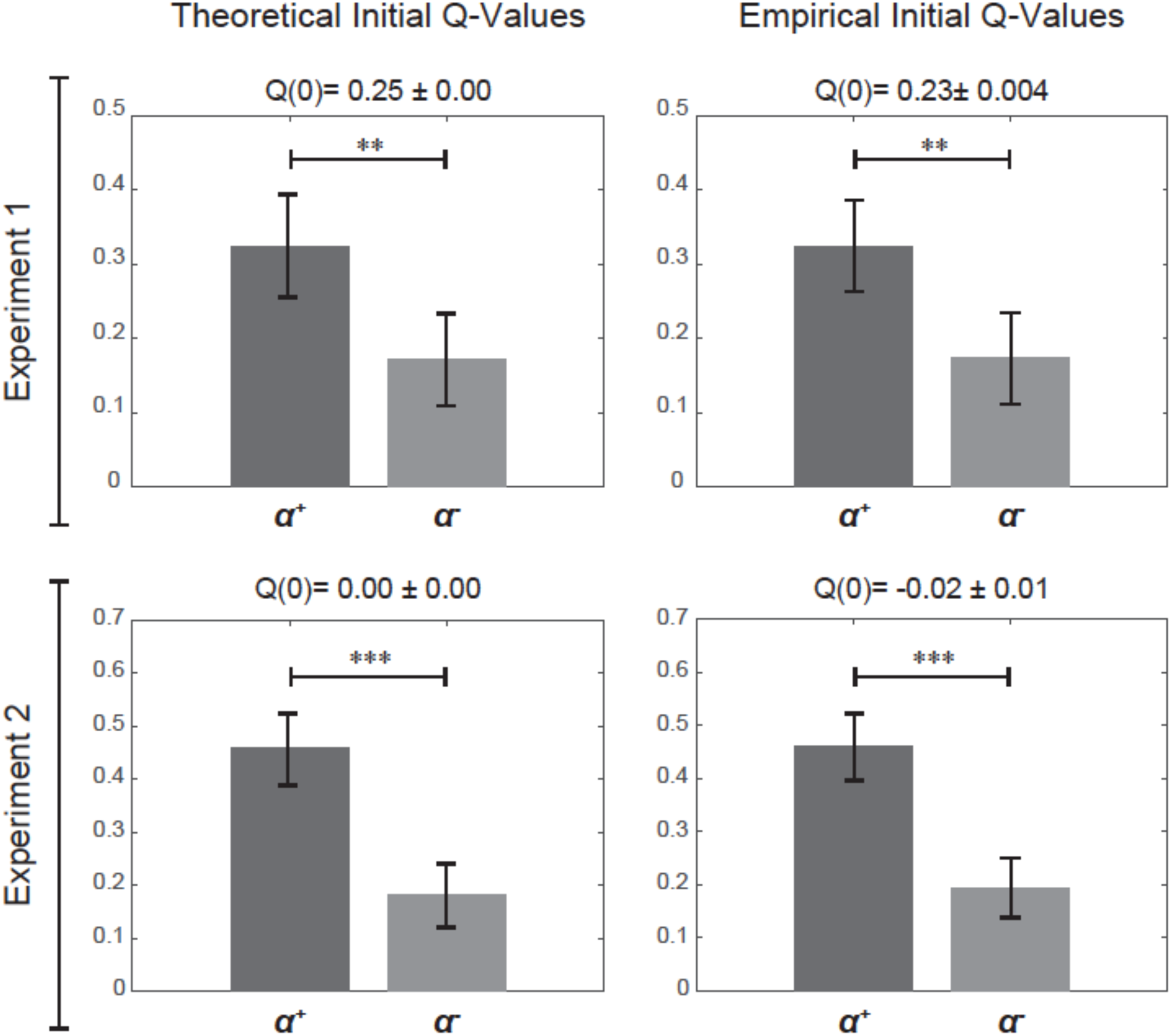
Bars represent for both experiments, learning rates retrieved assuming the initial Q-values equal to the mean between the best and the worst outcome *(leftmost panels)* or assuming the initial Q-values equal to the average outcome per symbol actually experienced during the training session *(rightmost panels)*. ***p<0.001 and **p<0.01, one-sample, two-tailed t-test.

### Conclusions

Supplementary analyses confirm the robustness of our results and the stable nature of optimistic reinforcement learning. Firstly, we fully replicated our behavioral and computational results in a second experiment including reward and actual monetary punishments (**Fig. S3**). We found that the learning asymmetry was robust to different settings and analyses (in all learning phases and in all contingency type; **Fig. S4**). Finally, the results were robust using initial Q-values empirically derived from the training session. This robustness of the optimistic reinforcement learning to a variety of situations corroborates our conclusions, placing the good news/bad news effect on the top of a low reinforcement learning bias.

### Supplementary Methods: model recovery

In order to verify that the parameters optimization procedure did not introduce systematic biases in the parameters’ value and to verify that both learning rate asymmetry and the exploitative behavior can be independently detected by our task and model, we run additional model simulations. We simulated four different types of subjects (N=1000 virtual subjects per computational phenotype): RW subjects (symmetric learning rates and higher exploration rate), RW± subjects (asymmetric learning rates and lower exploration rate), RW-exploitative subjects (symmetric learning rates and lower exploration rate), and RW±-explorative subjects (asymmetric learning rates and higher exploration rate). So basically our models simulations included the two computational phenotypes observed in our data plus two additional “hybrid” phenotypes. The results of the parameters optimizations indicated that the computational characteristics of each group were retrieved correctly (**Fig. S7**). Thus, the learning asymmetry and the tendency to exploit have to be considered as two independent features associated with optimistic behavior and not as an artifact of the model optimization procedure.

**Fig. S7:**
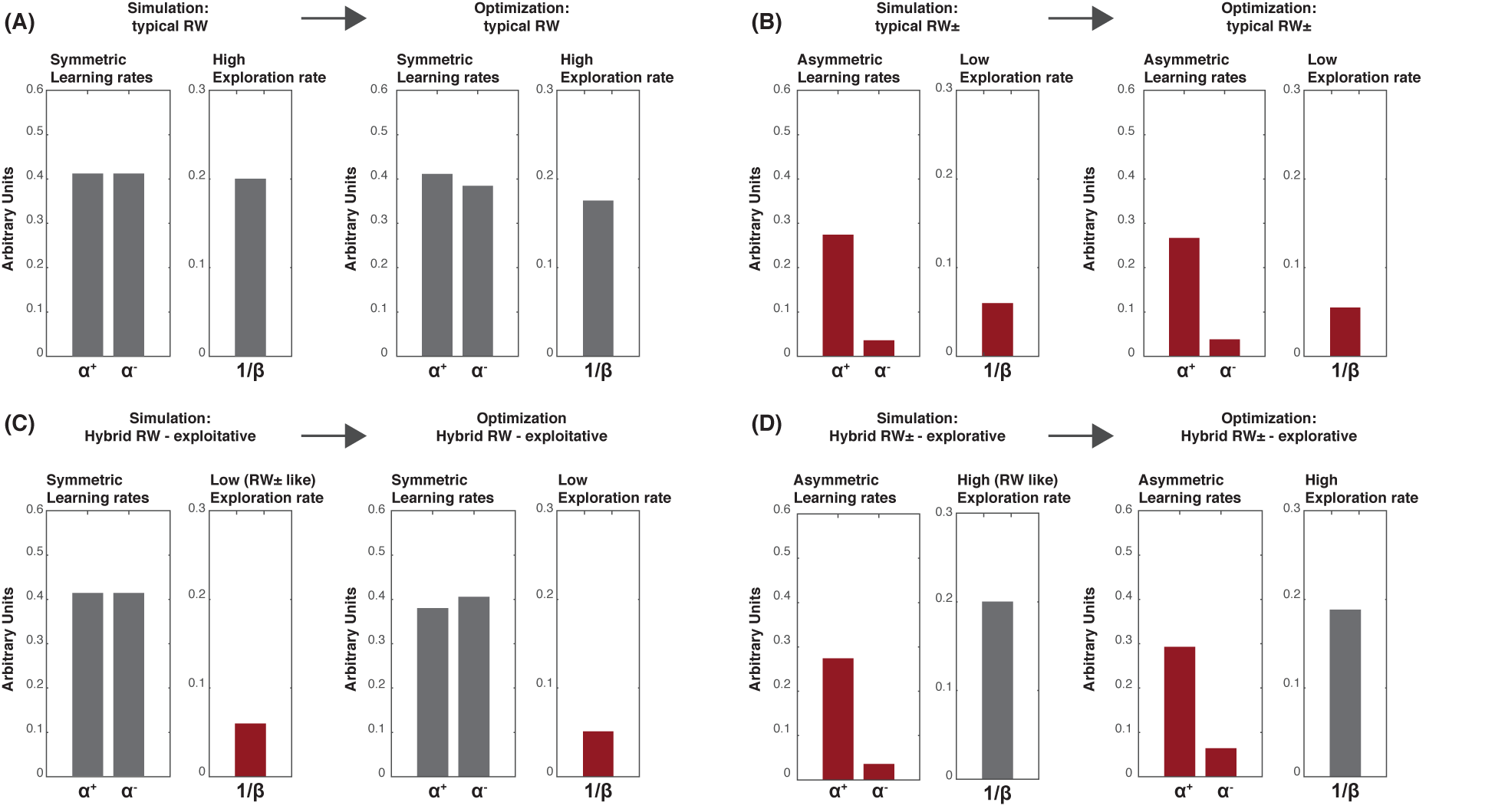
validation of the model optimization procedure. In each panel, the histograms represent the median parameters value used in the simulations (in the leftmost side of each panel: “Simulation”) and the parameter retrieved using the same method used for the behavioral data (in the rightmost side of each panel: “Optimization”). (**A**) Typical RW subjects (symmetric learning rates and higher temperature). (**B**) Typical RW± subjects (asymmetric learning rates and lower temperature). (**C**) Hybrid RW-exploitative subjects (symmetric learning rates and lower temperature). (**D**) Hybrid RW±-explorative subjects (asymmetric learning rates and higher temperature).

## References

1. Weinstein, N. D. Unrealistic Optimism About Future Life events. J. Pers. Soc. Psychol. 39, 806–820 (1980).

2. Shepperd, J. a., Klein, W. M. P., Waters, E. a. & Weinstein, N. D. Taking Stock of Unrealistic Optimism. Perspect. Psychol. Sci. 8, 395–411 (2013).

3. Shepperd, J. A., Waters, E. A., Weinstein, N. D. & Klein, W. M. P. A Primer on Unrealistic Optimism. Curr. Dir. Psychol. Sci. 24, 232–237 (2015).

4. Shepperd, J. a., Ouellette, J. a. & Fernandez, J. K. Abandoning unrealistic optimism: Performance estimates and the temporal proximity of self-relevant feedback. J. Pers. Soc. Psychol. 70, 844–855 (1996).

5. Waters, E. A. et al. Correlates of unrealistic risk beliefs in a nationally representative sample. J Behav Med 34, 225–235 (2011).

6. Schoenbaum, M. Do smokers understand the mortality effects of smoking? Evidence from the health and retirement survey. Am. J. Public Health 87, 755–759 (1997).

7. Sharot, T., Korn, C. W. & Dolan, R. J. How unrealistic optimism is maintained in the face of reality. Nat. Neurosci. 14, 1475–1479 (2011).

8. Eil, D. & Rao, J. M. The good news-bad news effect: Asymmetric processing of objective information about yourself. Am. Econ. J. Microeconomics 3, 114–138 (2011).

9. Sharot, T. & Garrett, N. Forming Beliefs: Why Valence Matters. Trends Cogn. Sci. 20, 25–33 (2016).

10. Sharot, T., Riccardi, A. M., Raio, C. M. & Phelps, E. A. Neural mechanisms mediating optimism bias. Nature 450, 102–105 (2007).

11. Moutsiana, C. et al. Human development of the ability to learn from bad news. Proc. Natl. Acad. Sci. 110, 16396–16401 (2013).

12. Garrett, N. et al. Losing the rose tinted glasses: neural substrates of unbiased belief updating in depression. Front. Hum. Neurosci. 8, 639 (2014).

13. Moutsiana, C., Charpentier, C. J., Garrett, N., Cohen, M. X. & Sharot, T. Human Frontal-Subcortical Circuit and Asymmetric Belief Updating. J. Neurosci. 35, 14077–14085 (2015).

14. Garrison, J., Erdeniz, B. & Done, J. Prediction error in reinforcement learning: A meta-analysis of neuroimaging studies. Neurosci. Biobehav. Rev. 37, 1297–1310 (2013).

15. Worbe, Y. et al. Reinforcement Learning and Gilles de la Tourette Syndrome. Arch. Gen. Psychiatry 68, 1257–1266 (2011).

16. Palminteri, S., Boraud, T., Lafargue, G., Dubois, B. & Pessiglione, M. Brain Hemispheres Selectively Track the Expected Value of Contralateral Options. J. Neurosci. 29, 13465–13472 (2009).

17. Palminteri, S. et al. Critical roles for anterior insula and dorsal striatum in punishment-based avoidance learning. Neuron 76, 998–1009 (2012).

18. Sutton, R. S. & Barto, A. G. Introduction to Reinforcement Learning. MIT Press Cambridge 135, (1998).

19. Rescorla, R. A. & Wagner, A. R. in Classical conditioning: current research and theory 64–99 (Appleton Century Crofts, 1972).

20. Daunizeau, J., Adam, V. & Rigoux, L. VBA : A Probabilistic Treatment of Nonlinear Models for Neurobiological and Behavioural Data. PLoS Comput. Biol. 10, e1003441 (2014).

21. O’Doherty, J. P., Hampton, A. & Kim, H. Model-based fMRI and its application to reward learning and decision making. Ann. N. Y. Acad. Sci. 1104, 35–53 (2007).

22. Shah, P., Harris, A. J. L., Bird, G., Catmur, C. & Hahn, U. A pessimistic view of optimistic belief updating. Cogn. Psychol. (2016). doi:10.1016/j.cogpsych.2016.05.004

23. Sharot, T., Garrett, N. & Psychology, E. The Myth of a Pessimistic View of Optimistic Belief Updating – A Commentary on Shah et al. Available SSRN 2811752 (2016).

24. Doll, B. B., Hutchison, K. E. & Frank, M. J. Dopaminergic Genes Predict Individual Differences in Susceptibility to Confirmation Bias. J. Neurosci. 31, 6188–6198 (2011).

25. Niv, Y. et al. Reinforcement Learning in Multidimensional Environments Relies on Attention Mechanisms. J. Neurosci. 35, 8145–8157 (2015).

26. Kahneman, D. & Tversky, A. Prospect Theory: An Analysis of Decision under Risk. Econometrica 47, 263–292 (1979).

27. Huys, Q. J. M., Maia, T. V & Frank, M. J. Computational psychiatry as a bridge from neuroscience to clinical applications. Nat. Neurosci. 19, 404–13 (2016).

28. Sharot, T. The Optimism Bias: Why We’re Wired to Look on the Bright Side. (Robinson, 2012).

29. Gifford, R. The dragons of inaction: pychological barriers that limit climate change mitigation and adaptation. Am. Psychol. 66, 290–302 (2011).

30. Sharot, T., Guitart-Masip, M., Korn, C. W., Chowdhury, R. & Dolan, R. J. How Dopamine Enhances an Optimism Bias in Humans. Curr. Biol. 22, 1477–1481 (2012).

31. Kuzmanovic, B., Jefferson, A. & Vogeley, K. The role of the neural reward circuitry in self-referential optimistic belief updates. Neuroimage 133, 151–162 (2016).

32. Skvortsova, V., Palminteri, S. & Pessiglione, M. Learning To Minimize Efforts versus Maximizing Rewards: Computational Principles and Neural Correlates. J. Neurosci. 34, 15621–15630 (2014).

33. Bartra, O., McGuire, J. T. & Kable, J. W. The valuation system: A coordinate-based meta-analysis of BOLD fMRI experiments examining neural correlates of subjective value. Neuroimage 76, 412–427 (2013).

34. Domenech, P. & Koechlin, E. Executive control and decision-making in the prefrontal cortex. Curr. Opin. Behav. Sci. 1, 101–106 (2015).

35. Kolling, N., Behrens, T. E. J., Wittmann, M. K. & Rushworth, M. F. S. Multiple signals in anterior cingulate cortex. Curr. Opin. Neurobiol. 37, 36–43 (2016).

36. Mathys, C., Daunizeau, J., Friston, K. J. & Stephan, K. E. A bayesian foundation for individual learning under uncertainty. Front. Hum. Neurosci. 5, 39 (2011).

37. Lebreton, M., Abitbol, R., Daunizeau, J. & Pessiglione, M. Automatic integration of confidence in the brain valuation signal. Nat. Neurosci. 18, 1159–1167 (2015).

38. Hampton, A. N., Bossaerts, P. & O’Doherty, J. P. The role of the ventromedial prefrontal cortex in abstract state-based inference during decision making in humans. J. Neurosci. 26, 8360–8367 (2006).

39. Van Den Bos, W., Cohen, M. X., Kahnt, T. & Crone, E. A. Striatum-medial prefrontal cortex connectivity predicts developmental changes in reinforcement learning. Cereb. Cortex 22, 1247–1255 (2012).

40. Frank, M. J., Moustafa, A. a, Haughey, H. M., Curran, T. & Hutchison, K. E. Genetic triple dissociation reveals multiple roles for dopamine in reinforcement learning. Proc. Natl. Acad. Sci. U. S. A. 104, 16311–16316 (2007).

41. Niv, Y., Edlund, J. A., Dayan, P. & O’Doherty, J. P. Neural prediction errors reveal a risk-sensitive reinforcement-learning process in the human brain. J. Neurosci. 32, 551–62 (2012).

42. Carver, C. S., Scheier, M. F. & Segerstrom, S. C. Optimism. Clin. Psychol. Rev. 30, 879–889 (2010).

43. Tindle, H. A. et al. Optimism, Cynical Hostility, and Incident Coronary Heart Disease and Mortality in the Women’s Health Initiative. Circulation 120, 656–662 (2009).

44. Macleod, A. K. & Conway, C. Well-being and the anticipation of future positive experiences: The role of income, social networks, and planning ability. Cogn. Emot. 19, 357–74 (2005).

45. Johnson, D. D. P. & Fowler, J. H. The evolution of overconfidence. Nature 477, 317–320 (2011).

46. Cazé, R. D. & van der Meer, M. A. A. Adaptive properties of differential learning rates for positive and negative outcomes. Biol. Cybern. 107, 711–719 (2013).

47. Raafat, R. M., Chater, N. & Frith, C. Herding in humans. Trends Cogn. Sci. 13, 420–428 (2009).

48. Hills, T. T., Todd, P. M., Lazer, D., Redish, A. D. & Couzin, I. D. Exploration versus exploitation in space, mind, and society. Trends Cogn. Sci. 19, 46–54 (2015).

49. Palminteri, S., Khamassi, M., Joffily, M. & Coricelli, G. Contextual modulation of value signals in reward and punishment learning. Nat. Commun. 6, 8096 (2015).

50. Popper, K. The logic of scientific discovery. Routledge (2005).

51. Dienes, Z. Understanding psychology as a science: An introduction to scientific and statistical inference. (Palgrave Macmillan, 2008).

52. Lebreton, M. & Palminteri, S. Assessing inter-individual variability in brain-behavior relationship with functional neuroimaging. bioRxiv 1–14 (2016). doi:http://dx.doi.org/10.1101/036772

## Supplementary References

1. Behrens, T. E. J., Woolrich, M. W., Walton, M. E. & Rushworth, M. F. S. Learning the value of information in an uncertain world. Nat. Neurosci. 10, 1214–1221 (2007).

2. Vinckier, F. et al. Confidence and psychosis: a neuro-computational account of contingency learning disruption by NMDA blockade. Mol. Psychiatry 1–10 (2015). doi:10.1038/mp.2015.73

3. Cazé, R. D. & van der Meer, M. A. A. Adaptive properties of differential learning rates for positive and negative outcomes. Biol. Cybern. 107, 711–719 (2013).

